# Vitamin D receptor cistrome-transcriptome analyses establishes quantitatively distinct receptor genomic interactions in African American prostate cancer regulated by BAZ1A

**DOI:** 10.1101/2022.01.31.478573

**Authors:** Manjunath Siddappa, Shahid Hussain, Sajad A. Wani, Hancong Tang, Jaimie S. Gray, Hedieh Jafari, Hsuchang Wu, Mark D. Long, Isra Elhussin, Balasubramanyam Karanam, Honghe Wang, Rebecca Morgan, Gary Hardiman, Isaacson B. Adelani, Solomon O. Rotimi, Adam R Murphy, Larisa Nonn, Melissa B Davis, Rick A Kittles, Chanita Hughes Halbert, Lara E. Sucheston-Campbell, Clayton Yates, Moray J. Campbell

**Affiliations:** Pharmaceutics and Pharmaceutical Chemistry, College of Pharmacy, The Ohio State University, Columbus, OH 43210; Department of Biostatistics and Bioinformatics, Roswell Park Comprehensive Cancer Center, Buffalo, NY 14263; Department of Biology and Center for Cancer Research, Tuskegee University, Tuskegee, AL 36088; School of Biological Sciences, Institute for Global Food Security, Queen’s University Belfast, Belfast, United Kingdom; Department of Medicine, Medical University of South Carolina, Charleston, SC 29425; Department of Biochemistry, Covenant University, KM. 10 Idiroko Road, Canaan Land, 112104, Ota, Ogun State, Nigeria; Department of Urology, Northwestern Medicine, Chicago, Illinois; Department of Pathology, University of Illinois at Chicago, Chicago, Illinois, USA; Department of Surgery, Weill Cornell Medicine, 420 E 70th Street, New York City, NY, 10021, USA.; Division of Health Equities, Department of Population Sciences, City of Hope, Duarte, 91010, CA, USA; Department of Population and Public Health Sciences, University of Southern California, Los Angeles, CA 30089; Norris Comprehensive Cancer Center, University of Southern California, Los Angeles, CA 20089; Division of Pharmacy Practice and Science, College of Pharmacy, The Ohio State University, Columbus, OH 43210.; Department of Veterinary Biosciences, College of Veterinary Medicine, The Ohio State University, Columbus, OH 43210

**Author notes:** These authors contributed equally to the study (Manjunath Siddappa, Shahid Hussain). Dept. of Veterinary Services and Animal Husbandry, Karnataka, India, 580001 (Manjunath Siddappa). **Author emails** Manjunath Siddappa; Shahid Hussain Sofi; Sajad A. Wani; Hancong Tang; Hedieh Jafari; Jaimie S. Gray; Hsuchang Wu; Mark D. Long; Isra Elhussin; Balasubramanyam Karanam; Honghe Wang; Rebecca Morgan; Gary Hardiman; Isaacson B. Adelani; Solomon O. Rotimi; Adam R Murphy; Larisa Nonn; Melissa B Davis; Rick A Kittles; Chanita Hughes-Halbert; Lara E. Sucheston-Campbell; Clayton Yates.

**Keywords:** Vitamin D Receptor, Prostate Cancer, Health Disparities, cistrome, transcriptome, SWI/SNF (SWItch/Sucrose Non-Fermentable), BAZ1A, SMARCA5

## Abstract

**Background:** African American (AA) prostate cancer (PCa) appears uniquely sensitive to 1α,25(OH)_2_D_3_ signaling, compared to European American (EA) PCa, but the extent and impact of vitamin D receptor genomic functions remain poorly defined.

**Results:** A panel of EA and AA prostate epithelial cells (EA: HPr1-AR, LNCaP, AA: RC43N, RC43T, RC77N, RC77T) were analyzed with RIME to reveal the cell-specific composition of the VDR- complex. 1α,25(OH)_2_D_3_-dependent ATAC-Seq revealed the greatest impact on nucleosome positioning in RC43N and RC43T, with gain of nucleosome-free at enhancer regions. VDR ChIP-Seq identified stronger and more frequent VDR binding in RC43N and RC43T that was enriched for a larger and distinct motif repertoire, than EA cells. VDR binding significantly overlapped with core circadian rhythm transcription factors in AA cell line models. RNA-Seq also revealed significantly stronger 1α,25(OH)_2_D_3_ dependent VDR transcriptional responses enriched for circadian rhythm and inflammation networks in AA cells. Whilst RC43N was most responsive, RC43T displayed distorted responses. Significantly reduced *BAZ1A/SMARCA5* in AA PCa samples was identified, and restored BAZ1A expression uniquely and significantly increased 1*α*,25(OH)_2_D_3_-regulated VDR targets in AA cells. These VDR-dependent cistrome-annotated genes were also uniquely and most significantly identified in three cohorts of AA PCa patients.

**Conclusion:** These data suggest VDR transcriptional control in the prostate is more potent and dynamic in AA men, and primed to govern inflammatory and circadian pathways. Reduced BAZ1A/SMARCA5 expression and/or reduced environmentally-regulated serum vitamin D_3_ levels suppress these actions. Therefore, the VDR axis lies at the cross-roads of biopsychosocial processes including stress responses, access to quality early detection and treatment, social determinants and that collectively contribute to PCa health disparities.

## BACKGROUND

The challenges of accurate early diagnosis of aggressive prostate tumors are amplified by a significant and devastating racial health disparity facing African American (AA) prostate cancer (PCa) patients. PCa in AA men is more aggressive and occurs at a younger age than in European American (EA) counterparts[1]. Current understanding of the drivers of PCa health disparities leverages genetic [2–4] and epigenetic [5–10] factors, combined with a range of biopsychosocial processes. The combined interactions of these intrinsic and extrinsic factors appear to drive uniquely aggressive disease in AA men. That there are biological drivers of PCa health disparities is supported strongly by the increased incidence of *TMPRRS2* and *ETS* fusion genes, which can act as cancer-drivers[11, 12], in EA PCa patients compared to AA counterparts [8, 13, 14].

To identify biological drivers of AA PCa we have focused on the vitamin D receptor (VDR), as this nuclear hormone receptor is implicated in cancer health disparities[15–17]. Ultraviolet B (UVB) radiation in the skin converts 7-dehydrocholesterol to cholecalciferol which is metabolized further to active 1α,25(OH)_2_D_3_. However, UVB radiation also degrades folic acid, which is also essential, and during evolution and adaptation a strong inverse correlation between skin pigmentation and latitude has arisen [18, 19]. In more recent human history relatively rapid dispersion and lifestyle changes have resulted in many individuals living in an UVB environment which differ from those of their ancestors.

Significant associations have been observed between either altered VDR signaling or low serum vitamin D_3_ levels and cancer incidence and progression risks amongst AA PCa patients [15-17, 20-32]. Furthermore, vitamin D_3_ supplementation has been evaluated in large-scale randomized trials to define individual health benefits[33, 34]. In the VITAL cohort of approximately 26,000 men and women vitamin D_3_ supplementation had no overall impact on cancer incidence, although AA participants experienced a suggestive 23% (p=0.07) reduction in cancer risk, indicating that larger cohorts maybe more informative in terms of defining the benefits of supplementation for these men [35–37]. Furthermore links between VDR signaling and anti-inflammatory actions appear strongest in AA PCa patients compared to EA counterparts [26]. Specifically, vitamin D_3_ supplementation prior to radical prostatectomy significantly altered tumor expression of genes associated with inflammation in the prostates of AA compared to EA patients[38].

Aside from calcemic functions, the VDR regulates gene networks that control cell cycle progression, promote cell differentiation, and immunomodulatory events (reviewed in[39]). More recently, independently of genomic ancestry, VDR signaling in has been associated with regulation of the circadian rhythm[40], reflecting a wider function for nuclear receptors to regulate this key process (reviewed in[41]). Together these data suggest that AA men may have a unique vulnerability to inadequate VDR signaling, which was potentially overlooked in the clinical evaluation of vitamin D_3_ analogs in PCa cohorts that largely consisted of EA men[42, 43].

Therefore, we sought to define VDR signaling capacity in EA and AA prostate cell models and clinical cohorts. We utilized EA and AA prostate cell models with confirmed ancestry; namely non-malignant EA prostate cells (HPr1AR)[44, 45], EA PCa (LNCaP), non-malignant prostate cells AA (RC43N, RC77N) and AA PCa (RC43T, RC77T)[46, 47]. Using these models, we defined the basal and 1α,25(OH)_2_D_3_-regulated VDR protein interactome (RIME), the VDR cistrome (ChIP-Seq and ATAC-Seq) and the VDR transcriptome (RNA-Seq). These interactome-cistrome-transcriptome relationships were associated with outcomes in several clinical cohorts; 1) in AA and EA men with high grade prostatic intraepithelial neoplasia (HGPIN) who were at risk of progression to PCa; 2); a PCa chemoprevention trial where AA and EA men with localized PCa were supplemented with vitamin D_3_ and 3) a PCa cohort of AA and EA men with gene expression data, clinical data and measured serum vitamin D_3_ levels. These approaches combine to reveal that AA cells displayed more potent, and qualitatively and quantitatively different VDR signaling than EA cells, and reduced expression of in AA PCa of members of the SWI/SNF ATP-dependent chromatin remodeling complex, including *BAZ1A* and *SMARCA5* suppressed VDR-dependent gene expression.

Together these studies support the concept that AA prostate cells display greater VDR genomic signaling capacity than EA counterparts and that disruption through altered expression of the BAZ1A/SMARCA5 chromatin remodeling complex primes cells to lose the ability to control immunomodulatory and circadian signaling.

## RESULTS

### The composition of the VDR complex and response to 1α,25(OH)_2_D_3_ differs significantly across prostate epithelial cells

We measured the anti-proliferative responses towards 1α,25(OH)_2_D_3_ across the six EA and AA cell models. Using optimal seeding conditions all cells had approximately equal doubling times (data not shown). **Figure 1A** demonstrates that the cancer cell lines (LNCaP, RC43T and RC77T) had more potent anti-proliferative responses towards 1α,25(OH)_2_D_3_ than the non-malignant counterparts. Only RC43T and LNCaP cells form colonies and RC43N and HPr1AR do not. LNCaP displayed a clear 1α,25(OH)_2_D_3_-mediated inhibition of clonal proliferation, RC43T colony number was only stimulated by 1α,25(OH)_2_D_3_ treatment (**Figure 1B**).

**Figure 1.**
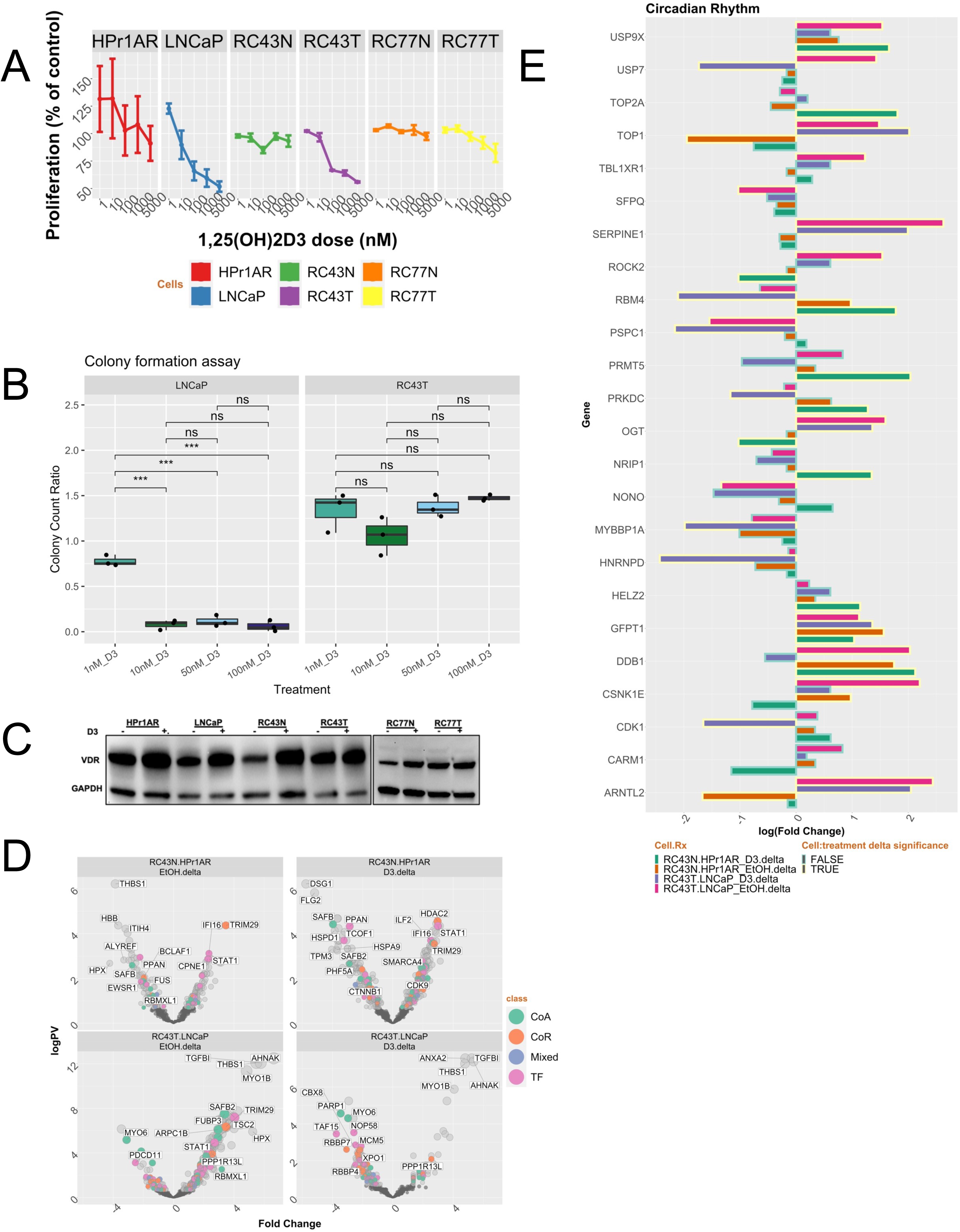
Expression of VDR and responses to 1α,25(OH)_2_D_3_ in African and European American Cell lines. **A.** The European American (EA) non-malignant HPr1AR and malignant LNCaP prostate cells and the African American (AA) non-malignant RC43N and RC77N, and malignant RC43T and RC77T cells were plated at optimal seeding density in 96 well white walled plates and grown for 96h with the indicated nM doses of 1α,25(OH)_2_D_3_ or vehicle control. Measurements of cellular levels of ATP, as an indicator of cell viability were performed. Each measurement was performed in biological triplicates and in triplicate wells. **B.** Clonal proliferation was measured after 10 days growth with the indicated doses of 1α,25(OH)_2_D_3_ or vehicle control. **C.** Western immunoblot measurements of VDR levels after 1α,25(OH)_2_D_3_ treatment (100 nM, 24h) or vehicle control. **D.** RIME analyses was undertaken for VDR in the indicated cells and significantly different proteins were identified using an edgeR workflow. Volcano plots depicting expression levels compared to IgG controls in basal or 1α,25(OH)_2_D_3_-stimulated conditions (100 nM, 4h). Significant (p.adj < .1) differentially and uniquely enriched proteins in each cell and treatment condition were classified either as a Coactivator (CoA), Corepressor (CoR), Mixed function coregulator (Mixed) or transcription factor (TF). **E.** Circadian rhythm regulators were selected and visualized with bars with a yellow border being significant.

Next, we measured how the VDR complex differed between the four cell models. VDR protein levels were generally elevated following 1α,25(OH)_2_D_3_ treatment, which was most pronounced in RC43N cells (**Figure 1C**). Subsequently, we measured VDR interacting proteins in the presence of chromatin in the basal and 1α,25(OH)_2_D_3_-stimulated states (100 nM, 4h) using RIME [48]. The number of positively enriched proteins in the basal state were; HPr1AR, n = 116; LNCaP, n = 135; RC43N, n = 115; RC43T, n = 245. Following addition of 1α,25(OH)_2_D_3_ in HPr1AR and LNCaP the number of positively enriched proteins remained broadly constant (n = 119 and n = 110, respectively). However, more proteins were enriched in RC43N (n = 198) and profoundly fewer were enriched in RC43T (n = **15) (**Supplementary Table 1, Supplementary Figure 1B).

The enriched protein data were classified as to whether they were Coactivators (CoA), corepressors (CoR), mixed function coregulators (Mixed), Transcription factors (TF) or other proteins. **Supplementary Table 1** indicates the number of proteins by class, and the most significant class member, and suggests both commonality and diversity in VDR interacting partners across cell lines. For example, across all cells the most enriched TF in either the basal or 1α,25(OH)_2_D_3_-treated state was either VDR or its heterodimer partners RXR*α* and RXR*β*, whereas the most enriched cofactors or other TFs varied by cell line. Other commonly enriched classes of proteins included heat shock proteins (e.g. HSP90AA1), RNA binding proteins (e.g. RBM12B), DNA repair enzymes (e.g. XRCC family members) and DNA helicases (e.g. DDX3X).

To measure the extent of the differences in the VDR complex we calculated the significant differences between enrichments in the basal and 1α,25(OH)_2_D_3_-stimulated states across cells. This revealed significant differences of proteins enriched in either the basal and 1α,25(OH)_2_D_3_-stimulated states comparing RC43N to HPr1AR, and RC43T to LNCaP (**Figure 1D**). For example, 1α,25(OH)_2_D_3_ treatment of RC43N cells enriched HDAC2, TRIM29 and SMARCA4 in the VDR complex.

From the differentially enriched proteins, we next filtered those that are unique to each cell and treatment. **Table 1** summarizes the number of significant and unique gain and loss of protein enrichment by class, and the most significant class member, which reemphasizes the diversity in VDR interacting partners across cell lines. For example, between RC43N and HPr1AR there were a large number of uniquely enriched VDR-associated proteins including in the basal state, the CoR TRIM29 and following 1α,25(OH)_2_D_3_ treatment the CoR NOC2L is reduced and various cofactors that promote transcription associate including the TF STAT1 and DNA topoisomerase TOP2A. By contrast, there were fewer unique differences between RC43T and LNCaP cells and 1α,25(OH)_2_D_3_ treatment only uniquely changed 2 proteins between RC43T and LNCaP cells including TGFB1. Finally, we filtered these to the most pronounced changes and grouped them by coactivator and corepressor complexes (**Supplementary Figure 2, left, right)**. Also, proteins associated with circadian rhythm were examined. Notably, components of the circadian rhythm control were differentially enriched across cells, with gains of in RC43T compared to LNCaP and loss in RC43N compared to HPr1AR cells, and a similar reciprocal enrichment with NONO in RC43N and loss in RC43T comparisons (**Figure 1E**).

**Table 1:**
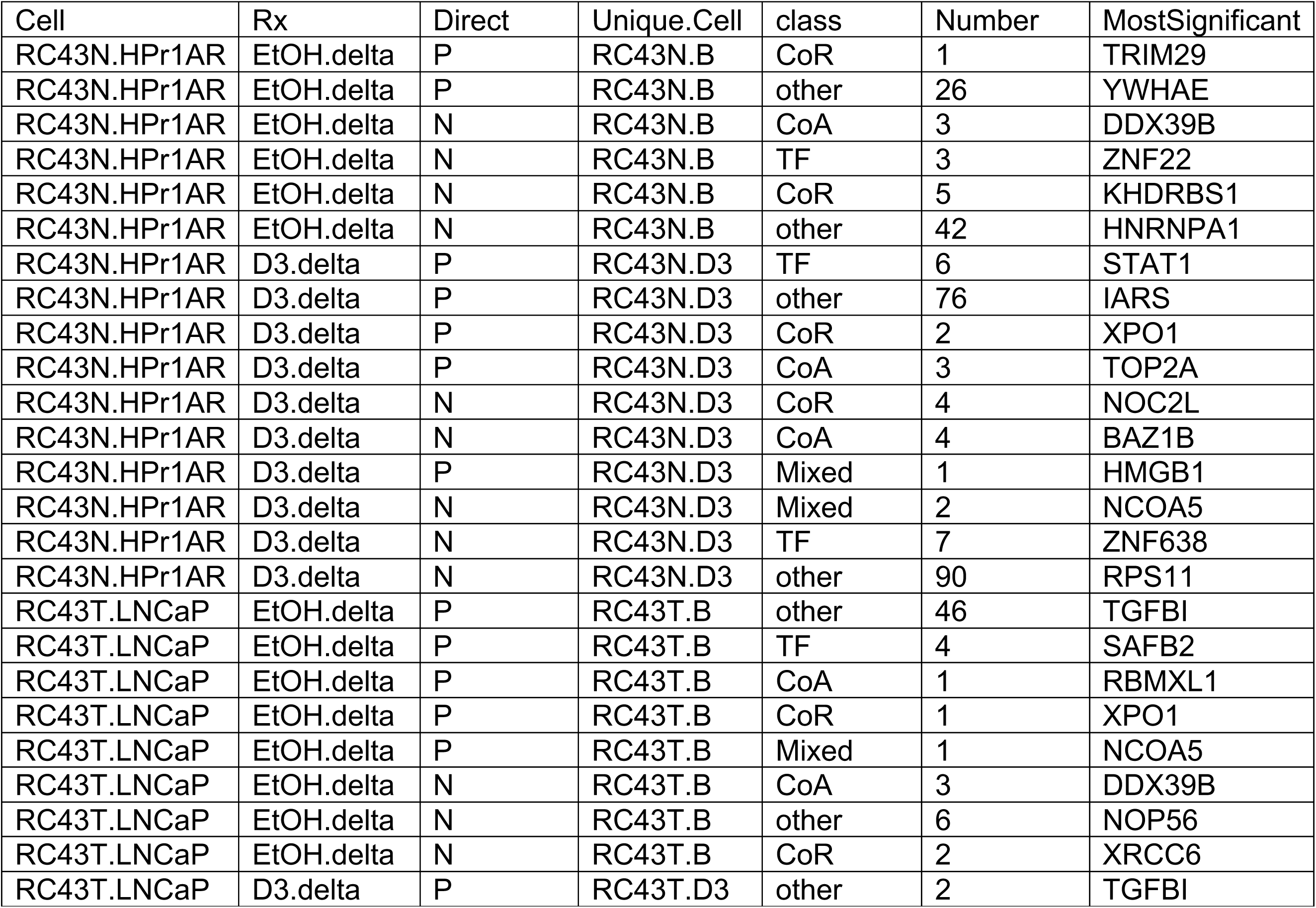
<u>Significantly and uniquely differentially enriched VDR-associated proteins</u>. RIME analyses captured VDR interacting proteins and significant differences were measured between cell types and in either the basal state of following 1α,25(OH)_2_D_3_ treatment (100 nM, 4h). Significant (p.adj < .1) differentially and uniquely enriched proteins in each cell and treatment condition were classified either as a Coactivator (CoA), Corepressor (CoR), Mixed function coregulator (Mixed) or transcription factor (TF), and the most significant member of each class in each condition is indicated.

### 1α,25(OH)_2_D_3_-dependent nucleosome positioning is distinct across cells

To understand the impact of the VDR complex on nucleosome positioning we undertook vehicle or 1α,25(OH)_2_D_3_ (100 nM, 4h) treatments and isolated chromatin for ATAC-Seq. Analyses of how 1α,25(OH)_2_D_3_ impacted the distribution of nucleosome-free (NF) mononucleosome (mono) (**Figure 2A, left, right**) and regions, revealed the greater 1α,25(OH)_2_D_3_-dependent impact in RC43N and RC43T than either HPr1AR or LNCaP. For example, in RC43N there were ∼11000 significant changes in 1α,25(OH)_2_D_3_-induced NF regions; 99.8% of which were gain in NF regions compared to the basal state. Similarly, 98% of the significant NF changes in HPr1AR were increases, but there were only ∼950 regions and a comparable response was also seen in LNCaP, although there were only ∼300 NF regions. There were no significant 1α,25(OH)_2_D_3_-regulated changes in mono regions in HPr1AR and RC43N, but there was a gain of ∼1800 mono regions in LNCaP cells. In RC43T, by contrast, 1α,25(OH)_2_D_3_-induced changes in NF and mono regions were predominantly a loss compared to controls. Thus, there were ∼ 1900 NF and ∼800 mono-nucleosome regions significantly impacted by 1α,25(OH)_2_D_3_ and these were mostly regions with a negative fold change compared to vehicle controls. Furthermore, the 1α,25(OH)_2_D_3_- regulated NF regions were significantly overlapped between RC43N and both RC43T and HPr1AR (p < 4.1e-230), whereas the LNCaP regions were distinct. Given that the shared NF regions between RC43N and RC43T are induced by 1α,25(OH)_2_D_3_ in RC43N but lost in RC43T, this further supports the concept that VDR function is distinct between these cells. Next, we analyzed how the NF and mono basal and 1α,25(OH)_2_D_3_-stimulated genomic partitions were distributed and enriched in prostate ChromHMM defined regions. The ChromHMM algorithm combines ChIP-Seq data to define epigenetic states and are publicly available for LNCaP cells[49]. The 1α,25(OH)_2_D_3_-stimulated NF and mono regions significantly enriched most frequently across Promoter regions (**Table 2**).

**Figure 2.**
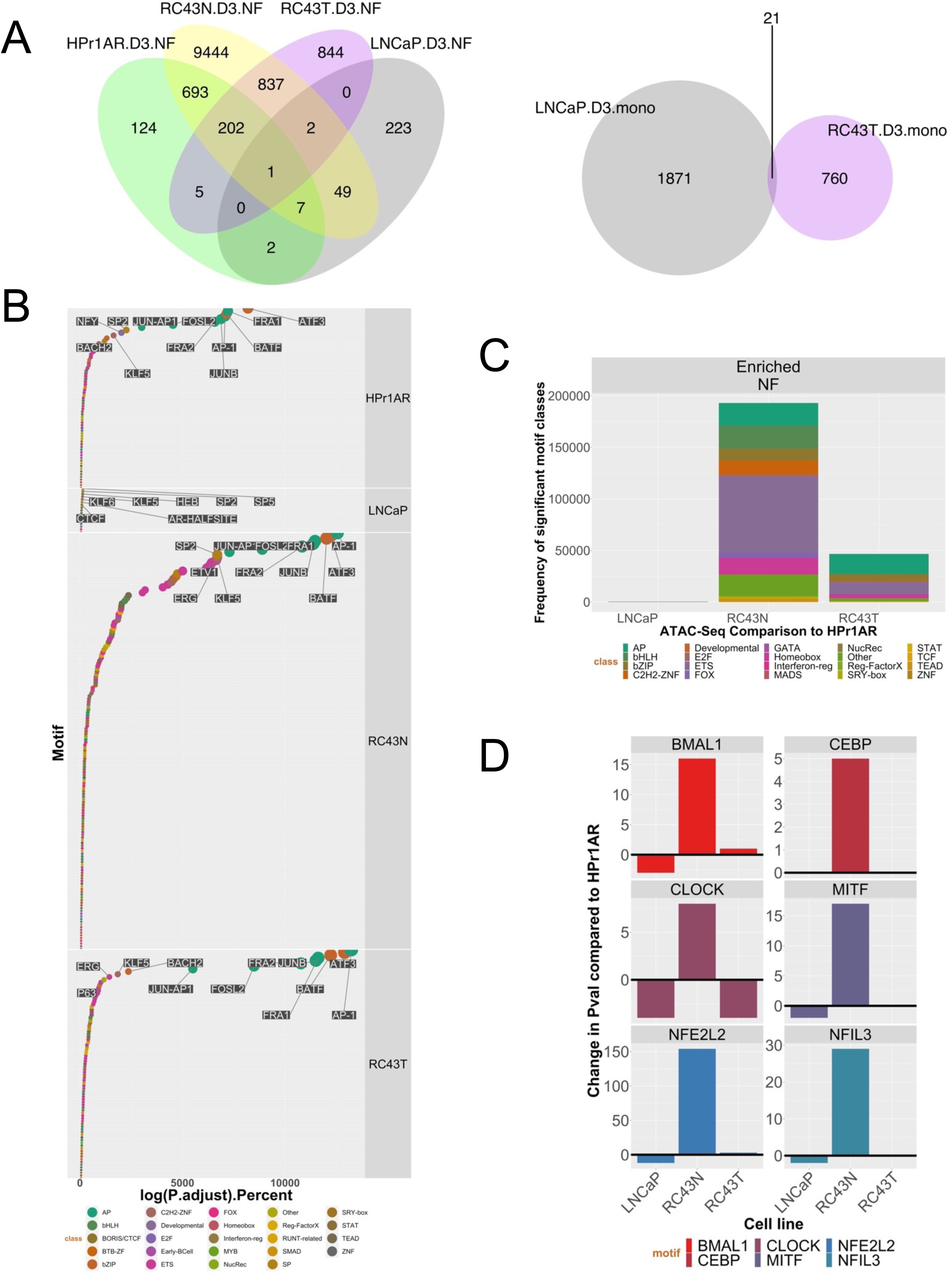
VDR ATAC-Seq in African and European American Cell lines. **A**. ATAC-Seq was undertaken in triplicate in HPr1AR, LNCaP, RC43N and RC43T following 1α,25(OH)_2_D_3_ treatment (100 nM, 4h) or vehicle control. FASTQ files were QC processed, aligned to hg38 (Rsubread), sorted and duplicates removed before further processing with ATACseqQC to generate nucleosome free and mono-nucleosome regions. Differential enrichment of regions was measured with csaw and the significantly different regions (p.adj < .1) were then intersected to generate the Venn diagrams of overlapping regions by a minimum of 1bp (ChIPpeakAnno). **B**. Motif enrichment in NF regions was undertaken with Homer and ranked by significance to visualize in descending significance. **C**. Frequency of enriched motifs by transcription factor families’ class. D. Enrichment of circadian rhythm transcription factors.

**Table 2:** <u>Enrichment of 1α,25(OH)_2_D_3_-induced nucleosome free (NF) and mononucleosome (mono) ChromHMM regions.</u> ATAC-Seq was undertaken following 1α,25(OH)_2_D_3_ treatment (100 nM, 4h) or vehicle control. Reads were processed by ATACseqQC to nucleosome fractions, and then differentially 1α,25(OH)_2_D_3_ enriched NF and mono free regions (p.adj < .1) were identified by csaw and were then overlapped with ChromHMM regions (ChromHMM) identified in LNCaP using bedtools and enrichment tested with a hypergeometric test (lower.tail = FALSE).

| Cell | RX | nuc | ChromHMM | logPval | Threshold | Direction |
| --- | --- | --- | --- | --- | --- | --- |
| LNCaP | D3 | mono | Promoter | 315.86 | Significant | Gain |
| RC43N | D3 | NF | Promoter | 315.86 | Significant | Gain |
| RC43T | D3 | mono | Promoter | 315.86 | Significant | Loss |
| HPr1AR | D3 | NF | Promoter | 315.51 | Significant | Gain |
| RC43T | D3 | NF | Promoter | 168.63 | Significant | Loss |
| LNCaP | D3 | NF | Promoter | 46.48 | Significant | Gain |
| RC43N | D3 | NF | Transcribed | 9.95 | Significant | Gain |
| RC43T | D3 | NF | Transcribed | 8.24 | Significant | Loss |
| RC43N | D3 | NF | Bivalent_Promoter | 1.40 | Significant | Gain |

We mined these NF and mono regions for motif enrichment to illuminate how changes in nucleosome positioning may impact transcription factor binding [50]. Consistent with the fact that the greatest number of 1α,25(OH)_2_D_3_-induced NF regions were identified RC43N cells, there were also more significant motif enrichments (by p value and percentage coverage) in this cell line. In **Figure 2B** and **Supplementary Figure 3A** the TF families are ranked by the significance of enrichment. For example, in RC43N, the most significantly enriched motifs included AP (e.g., AP-1), bZIP (e.g., ATF3), along with SP, FOX and ETS family members. This was broadly reflected by HPr1AR and RC43T cells, with RC43T cells showing a loss of potential TF interactions. LNCaP cells, by contrast, displayed fewer significant associations. Interestingly, one of the most enriched motifs in LNCaP was the AR/Half-site, which was also only enriched in RC43T regions. Enrichment in mono regions included SP (e.g., SP2), C2H2-ZNF (e.g. KLF5) and ETS (e.g. ELK4) family members ((**Figure 2B**; **Supplementary Figure 3A**).

We calculated delta values comparing the differences between cells in motif enrichment in the 1α,25(OH)_2_D_3_-induced NF regions (**Figure 2C**). Compared to HPr1AR cells, RC43N displayed the most striking gains in enrichment of multiple TF families including bHLH and FOX family members. The bHLH motifs included multiple TFs for circadian rhythm such as bMAL1 and CLOCK, which were either most clearly or exclusively enriched compared to other cells (**Figure 2D**). Similarly, FOX family members were highly enriched in RC43N, although it is notable that the FOXA1-AR motif enrichment is most prominent in LNCaP cells (**Supplementary Figure 3B**).

### The VDR cistrome is distinct across cells

The VDR cistrome was measured in cells treated with 1α,25(OH)_2_D_3_ (100 nM, 6h) and VDR peaks (FDR < 0.1) identified compared to IgG controls. Significant basal VDR binding was identified in LNCaP, RC43N and RC43T cells but not HPr1AR, and in all cells following 1α,25(OH)_2_D_3_-treatment (**Supplementary Table 2**). The basal peaks were highly overlapping (sharing at least 1 bp) with 305 shared sites across all three cells (**Figure 3A**). 1α,25(OH)_2_D_3_ treatment reduced the number of VDR binding sites in LNCaP and RC43T; there were 773 and 1560 basal VDR peaks in LNCaP and RC43T cells, which reduced to 322 and 30, respectively. Within each cell there was also a large overlap between basal and 1α,25(OH)_2_D_3_-stimulated sites (**Supplementary Figure 4A**) and the proximal/distal distribution of the VDR cistrome was broadly constant within a cell type regardless of 1α,25(OH)_2_D_3_ treatment (**Supplementary Figure 4B**).

**Figure 3.**
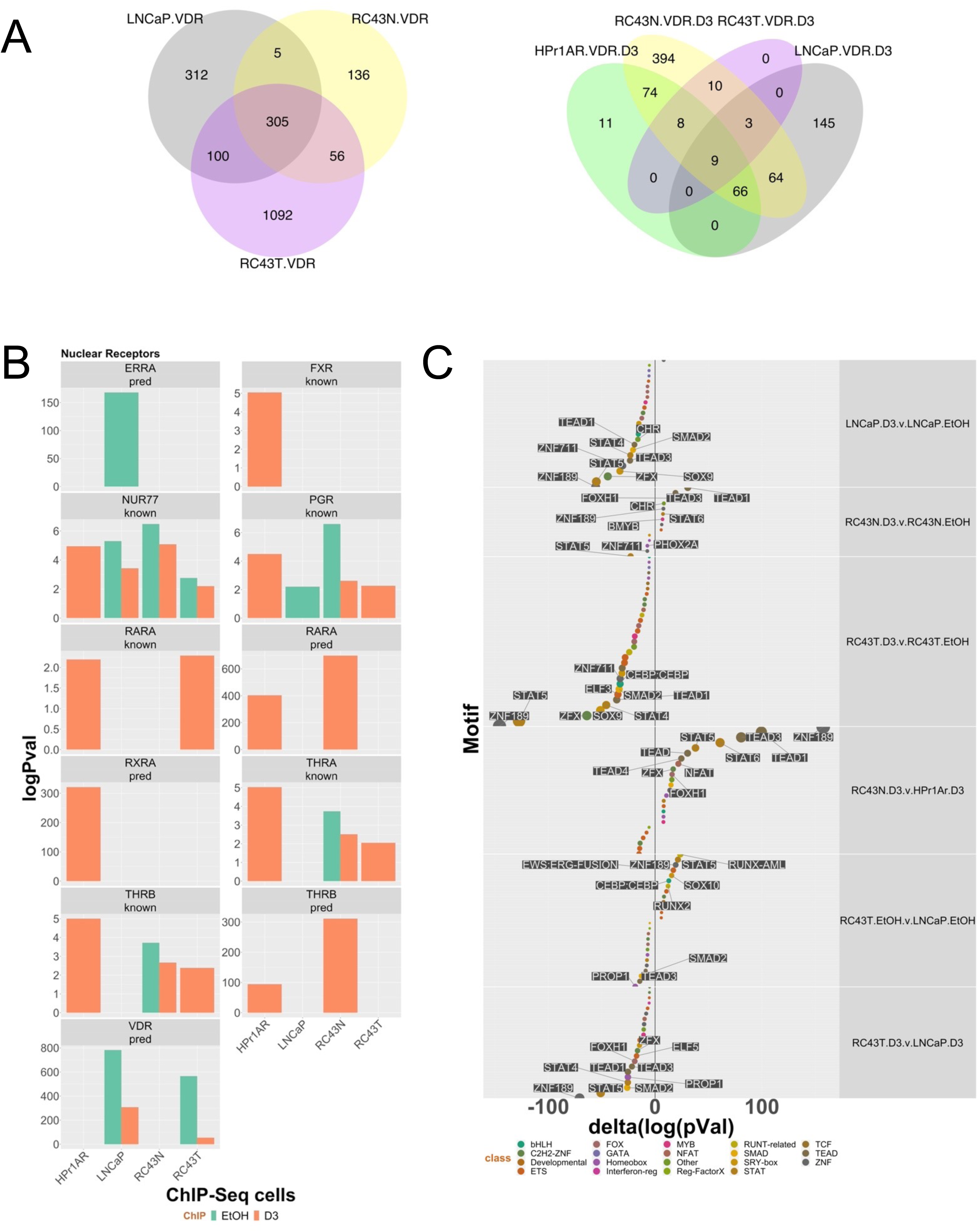
VDR ChIP-Seq in African and European American Cell lines. A. Basal and 1α,25(OH)_2_D_3_- stimulated (100 nM, 6h) VDR ChIP-Seq was undertaken in triplicate in HPr1AR, LNCaP, RC43N and RC43T. FASTQ files were QC processed, aligned to hg38 (Rsubread), sorted and duplicates removed before differential enrichment of regions was measured with csaw and the significantly different regions compared to IgG controls (p.adj < .1) were then intersected to generate the Venn diagrams of overlapping regions by a minimum of 1bp (ChIPpeakAnno). **B**. Significantly differentially enriched regions were identified by Homer and nuclear receptors are illustrated. C. Changes in motif enrichment were calculated (delta) and ranked by significance for visualization.

Similar to the ATAC-Seq data, we analyzed how the basal and 1α,25(OH)_2_D_3_-stimulated genomic partitions were distributed and enriched in prostate ChromHMM defined regions (**Table 3**). The basal VDR cistrome across cells was commonly enriched in Transcribed regions and Bivalent Promoters and remained so following 1α,25(OH)_2_D_3_ treatment. Other enrichments were cell specific, for example, Polycomb regions were only enriched in RC43T and LNCaP. Compared to the ATAC-Seq enrichments, which were frequent at Promoter regions, only basal VDR ChIP-Seq in LNCaP was enriched at Promoter regions, and only modestly so. Interestingly, this approach also revealed the significant impact of 1α,25(OH)_2_D_3_ treatment. For example, in LNCaP the log10(p.adj) of the basal VDR enrichment in Bivalent Promoters was 46, and this was reduced to 6 in the presence of 1α,25(OH)_2_D_3_, and similarly was 45 in RC43T in the basal state, and non-significant following 1α,25(OH)_2_D_3_ treatment. By contrast the score in RC43N was broadly equivalent in the basal and 1α,25(OH)_2_D_3_ states.

**Table 3:** <u>Enrichment of VDR ChIP-Seq and ChromHMM regions.</u> VDR ChIP-Seq was undertaken following 1α,25(OH)_2_D_3_ treatment (100 nM, 4h) or vehicle control. Differential VDR binding sites compared to IgG controls (p.adj < .1) were identified by csaw and were then overlapped with ChromHMM regions (ChromHMM) identified in LNCaP using bedtools and enrichment tested with a hypergeometric test (lower.tail = FALSE).

| Cell.ChIP | ChromHMM | logPV | Threshold |
| --- | --- | --- | --- |
| LNCaP.VDR | Bivalent_Promoter | 46.16 | Significant |
| RC43T.VDR | Transcribed | 45.74 | Significant |
| RC43T.VDR | Bivalent_Promoter | 40.84 | Significant |
| RC43T.VDR | Polycomb | 35.42 | Significant |
| LNCaP.VDR | Polycomb | 30.69 | Significant |
| LNCaP.VDR | Transcribed | 22.56 | Significant |
| RC43N.VDR.D3 | Bivalent_Promoter | 20.51 | Significant |
| RC43N.VDR | Bivalent_Promoter | 15.49 | Significant |
| LNCaP.VDR.D3 | Bivalent_Promoter | 6.37 | Significant |
| HPr1AR.VDR.D3 | Bivalent_Promoter | 4.63 | Significant |
| HPr1AR.VDR.D3 | Transcribed | 4.48 | Significant |
| LNCaP.VDR | Poised_Enhancer | 4.41 | Significant |
| RC43N.VDR.D3 | Transcribed | 3.98 | Significant |
| LNCaP.VDR.D3 | Transcribed | 3.07 | Significant |
| RC43N.VDR | Transcribed | 2.09 | Significant |
| LNCaP.VDR | Promoter | 1.56 | Significant |
| RC43T.VDR | Poised_Enhancer | 1.34 | Significant |

Motif analyses categorized significantly enrichment of TF families across cells and treatments (**Table 4)**. For example, the predicted VDR motif and the most significant nuclear receptor motif in LNCaP was also identified in RC43T. By contrast, the RAR*α* motif was the most significant nuclear receptor motif in HPr1AR and RC43N (**Figure 3B**). By contrast to the ATAC-Seq data, the enriched bHLH motifs did not include CLOCK, BMAL1, MITF and NFEs, but did include CEBP in all conditions. To gain further insight into how these motif enrichments changed by 1α,25(OH)_2_D_3_ treatment and across cell types, we calculated the delta log(p.adj) between groups, and visualized the most altered motifs (**Figure 3C**). This revealed, for example, in LNCaP, RC43N and RC43T cells that 1α,25(OH)_2_D_3_ treatment reduced enrichment for various motifs including STAT5. In fact, only in RC43N cells did 1α,25(OH)_2_D_3_ treatment increase motif enrichment notably for TEAD1 and TEAD3. Similarly, the circadian rhythm associated TF, NFAT, was more enriched in 1α,25(OH)_2_D_3_ treated RC43N than HPr1AR.

**Table 4:**
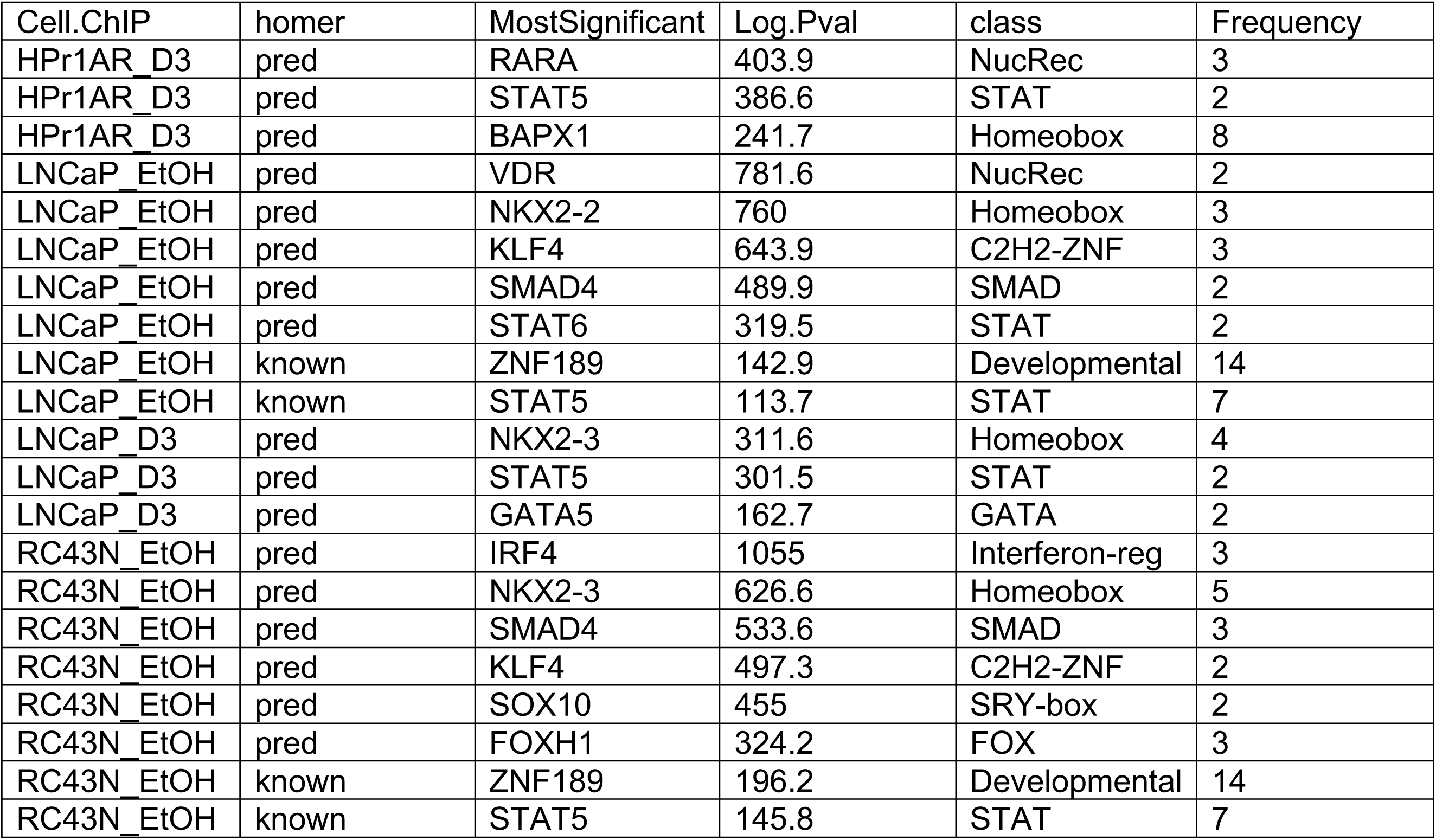
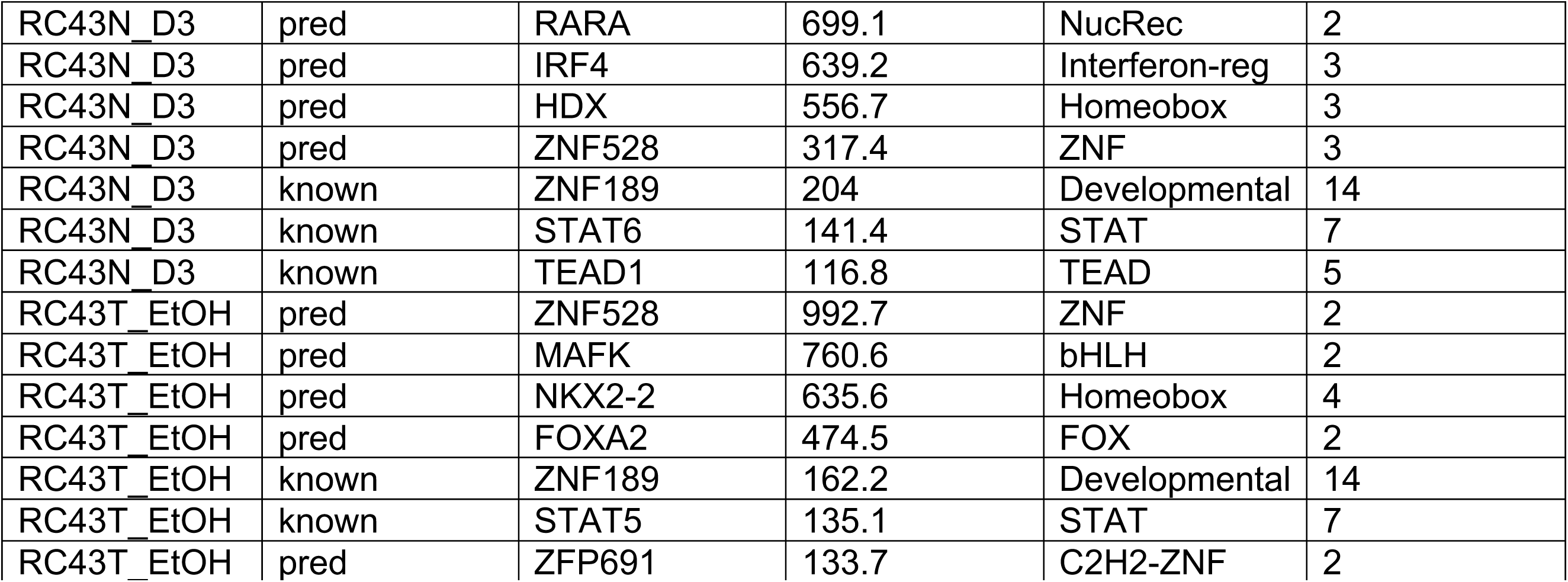
<u>Summary of motif enrichment by transcription family in the VDR dependent cistromes.</u> Significant VDR binding sites were analyzed by Homer to identify enrichment of motifs, by transcription factor family enrichment. Within each cistrome the known and predicted (pred) motifs are arranged by -log(p.adj) enrichment value for the most significant family member.

Next, VDR cistromes were analyzed for significant overlaps with TF and histone modifications contained in the CistromeDB collection (> 10,000 total ChIP-seq datasets, across > 1100 factors) using GIGGLE[51], which searches publicly available datasets within the Cistrome database and returns samples which contains peak sites co-incident with user-supplied data (**Supplementary Figure 5**); from these data we calculated the comboScore.adj as the negative log10 of the fisher.pval multiplied by the log2 of the odds.ratio. This approach reflected the motif analyses and provided additional strong evidence for VDR binding being co-incident with multiple TFs and histone modifications, and that the 1α,25(OH)_2_D_3_ treatment dynamically altered many of these relationships. These analyses also revealed clear changes in numerous ZNF transcription factors (magenta) and a clear under-enrichment of bHLH members in the basal RC43N cells (**Supplementary Figure 5A**). There were significant overlaps with nuclear receptor ChIP-Seq data, including for PPARs, RARs, and VDR, which were all most significant in RC43N (**Supplementary Figure 5B**). For example, a combined score for overlapping with VDR ChIP-Seq (calculated as the odds ratio multiplied by the -log10 (Fisher.p.adj)), was 1111 and 1285 for RC43N in the basal and 1α,25(OH)_2_D_3_-treated cells in non-malignant prostate epithelial cells. These were much reduced in HPr1AR and LNCaP (4 and 71, respectively). Similarly, there were 1α,25(OH)_2_D_3_-dependent increases in enrichment for other nuclear receptors including AR, ERα, and RAR*γ*. Analyzing a canonical list of 32 transcriptional regulators of circadian rhythm revealed they significantly overlapped with VDR cistromes most significantly in RC43N. The most frequent enrichments were in both the basal and 1α,25(OH)_2_D_3_-treated RC43N cells, and indeed were amongst the most significantly overlapping factors within any VDR cistrome, including NONO (comboScore.adj = 1745 RC43N 1α,25(OH)_2_D_3_-treated) (**Supplementary Figure 5C**).

Given that the median distance for enhancer to TSS is greater than 100 kb[52–54], the VDR cistromes (ATAC-Seq and ChIP-Seq) were related to genes within a 100 kb window up and downstream of genes. From these genes we measured which encoded for proteins that were also identified by RIME to interact with the VDR in either the basal or 1α,25(OH)_2_D_3_-stimulated state (**Table 5**). This suggested that ATAC-Seq data sets commonly annotated to genes that in many cases encoded for proteins that were positively enriched with the VDR. In HPr1AR, LNCaP and RC43N approximately equal numbers of basal and 1α,25(OH)_2_D_3_-stimulated proteins were enriched with VDR that were also targets of the cistrome data. However, this not the case in RC43T, which had a large number of overlaps between ATAC-Seq NF regions and the VDR interactome in the basal state but far fewer in the 1α,25(OH)_2_D_3_-stimulated state. Given the profound reduction of 1α,25(OH)_2_D_3_-stimulated NF regions in RC43T this may suggest a mechanism to limit the function of the VDR complex in this cell.

**Table 5:** <u>Genes annotated to 1α,25(OH)_2_D_3_-regulated nucleosome free regions and VDR binding sites overlap with VDR-interacting proteins.</u> Genes were annotated to ATAC-Seq or ChIP-Seq regions within 100kB and those genes overlapped with the positively enriched proteins within the same cell background. Enriched genes were classified either as a Coactivator (CoA), Corepressor (CoR), Mixed function coregulator (Mixed), transcription factor (TF) or other, and the most significant member of each class in each condition is indicated.

| Cell.cistrome | RIME.Rx | class | Number | MostSignificant |
| --- | --- | --- | --- | --- |
| HPr1AR.ATAC | D3 | other | 29 | ITGB4 |
| HPr1AR.ATAC | EtOH | other | 24 | YWHAQ |
| HPr1AR.ATAC | EtOH | TF | 4 | ARNTL2 |
| HPr1AR.ATAC | EtOH | CoA | 2 | RBMXL1 |
| HPr1AR.ATAC | D3 | TF | 2 | SSRP1 |
| HPr1AR.ATAC | EtOH | Mixed | 1 | CTBP2 |
| HPr1AR.ATAC | EtOH | CoR | 1 | HSPA8 |
| HPr1AR.ATAC | D3 | CoR | 1 | HSPA8 |
| HPr1AR.ATAC | D3 | CoA | 1 | SAFB |
| LNCaP.ATAC | EtOH | other | 3 | PGD |
| LNCaP.ATAC | D3 | other | 3 | PGD |
| LNCaP.ATAC | EtOH | TF | 1 | ENO1 |
| LNCaP.ATAC | EtOH | CoR | 1 | NOC2L |
| LNCaP.ATAC | D3 | TF | 1 | ENO1 |
| LNCaP.ATAC | D3 | CoR | 1 | NOC2L |
| RC43N.ATAC | D3 | other | 106 | CLTC |
| RC43N.ATAC | EtOH | other | 60 | CLTC |
| RC43N.ATAC | D3 | TF | 10 | VDR |
| RC43N.ATAC | D3 | CoA | 8 | PTBP1 |
| RC43N.ATAC | D3 | CoR | 7 | XRCC6 |
| RC43N.ATAC | EtOH | CoR | 4 | HSPA8 |
| RC43N.ATAC | EtOH | TF | 3 | VDR |
| RC43N.ATAC | EtOH | CoA | 2 | PTBP1 |
| RC43N.ATAC | D3 | Mixed | 1 | HMGB1 |
| RC43T.ATAC | EtOH | other | 158 | RBM12B |
| RC43T.ATAC | EtOH | TF | 17 | VDR |
| RC43T.ATAC | EtOH | CoA | 15 | SAFB |
| RC43T.ATAC | EtOH | CoR | 8 | XRCC5 |
| RC43T.ATAC | D3 | other | 8 | TGFBI |
| RC43T.ATAC | EtOH | Mixed | 2 | NCOA5 |
| RC43T.ATAC | D3 | TF | 1 | VDR |
| RC43T.ATAC | D3 | CoR | 1 | XRCC6 |
| LNCaP.ChIP | EtOH | other | 1 | TUBB4B |
| LNCaP.ChIP | EtOH | TF | 1 | GTF3C1 |
| LNCaP.ChIP | D3 | other | 1 | TUBB4B |
| RC43T.ChIP | EtOH | other | 7 | TNC |
| RC43T.ChIP | EtOH | CoR | 1 | TRIM29 |

### The VDR transcriptome is larger in RC43N and RC43T than HPr1AR and LNCaP cells

1α,25(OH)_2_D_3_-dependent transcriptomic analyses were undertaken in HPr1AR, LNCaP, RC43N and RC43T using both a standard and small RNA library preparation for RNA-Seq (100 nM, 8h). Similarity and principal component analyses revealed that experimental conditions explained the majority of variation in expression (**Supplementary Figure 6**). Reflecting the size of the VDR cistromes, the number of 1α,25(OH)_2_D_3_ regulated genes was larger in the AA than EA models. Differentially expressed genes in response to 1α,25(OH)_2_D_3_ (**Figure 4A**; DEGs; FDR < 0.1, FC > 1.3) were determined in HPr1AR (teal), LNCaP (orange), RC43N (indigo) and RC43T (pink). The top most regulated genes included the well-established VDR target gene CYP24A1; CYP24A1 was not in the top 5 labelled genes in RC43N, although was in top 20 DEGs. Genes that had a 1α,25(OH)_2_D_3_-dependent NF region (red symbol) and/or a VDR binding peak (symbol with dark border) in the same cell background were more apparent in AA cells.

**Figure 4.**
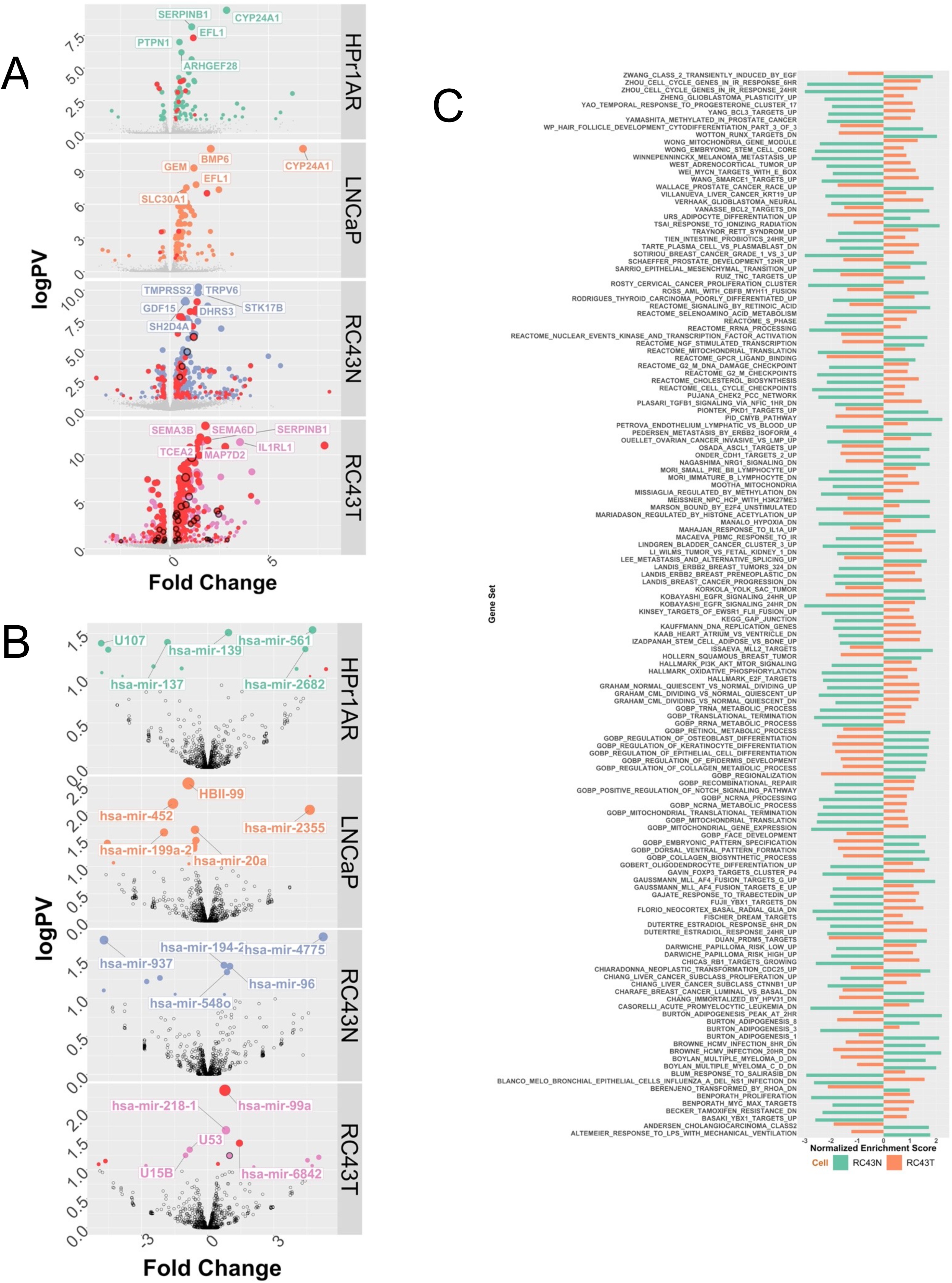
1α,25(OH)_2_D_3_-dependent RNA- and small RNA-Seq in African and European American Cell lines. **A**. **B**. Basal and 1α,25(OH)_2_D_3_-stimulated (100 nM, 8h) RNA-Seq was undertaken in triplicate in HPr1AR, LNCaP, RC43N and RC43T. FASTQ files were QC processed, aligned to hg38 (Rsubread), and processed with limma-voom and edgeR workflow to identify significant differentially enriched genes (DEGs) (logPV > 1 & absFC > .37) are illustrated on Volcano plots with top DEGs illustrated. C. GSEA analyses (Chemical Perturbations in GSEA) was undertaken and the terms identified and visualized where the normalized enrichment score was in the opposite direction in RC43N and RC43T.

This was also noticeable when considering the DEGs across cells of different genomic ancestry, namely the difference in expression between AA vs. EA tumors exposed to 1α,25(OH)_2_D_3_ (RC43T treated with 1α,25(OH)_2_D_3_ compared to LNCaP treated with 1α,25(OH)_2_D_3_) and difference in expression between AA vs. EA non-malignant cells exposed to 1α,25(OH)_2_D_3_ (RC43N treated with 1α,25(OH)_2_D_3_ compared to HPr1AR treated with 1α,25(OH)_2_D_3_). Small RNA expression patterns were comparable across cells in terms of magnitude and frequency with ∼20 short non-coding RNA, being regulated in each model, in a largely distinct manner (**Figure 4B**; DEGs; FDR < 0.1, FC > 1.3). 1α,25(OH)_2_D_3_-regulated miRNA that were also annotated to 1α,25(OH)_2_D_3_-dependent NF regions were identified in HPr1AR and RC43T and in the latter case included miR-99a.

We also used transcript-aware analyses to identify how 1α,25(OH)_2_D_3_-treatment induced differentially expressed transcripts (DETs). In each cell, 1α,25(OH)_2_D_3_-treatment induced significant DETs; HPr1AR, n=96; LNCaP, n=124; RC43N, n=136; RC43T, n=77, whereas the comparisons across cells were much larger (data not shown). Again, we categorized DETs functionally and in relationship to the cistrome date (**Table 6**). RC43N and RC43T cells displayed the highest proportion of DETs that were annotated to cistrome NF ATAC-Seq annotated genes. (RC43N, 84 out of 136; RC43T, 71 out of 77). Of these, 10 were TFs, and the most significant was ZNF83. Similarly, the coactivator SMARCA4 was an NF ATAC-Seq annotated gene and was a significant DET.

**Table 6:**
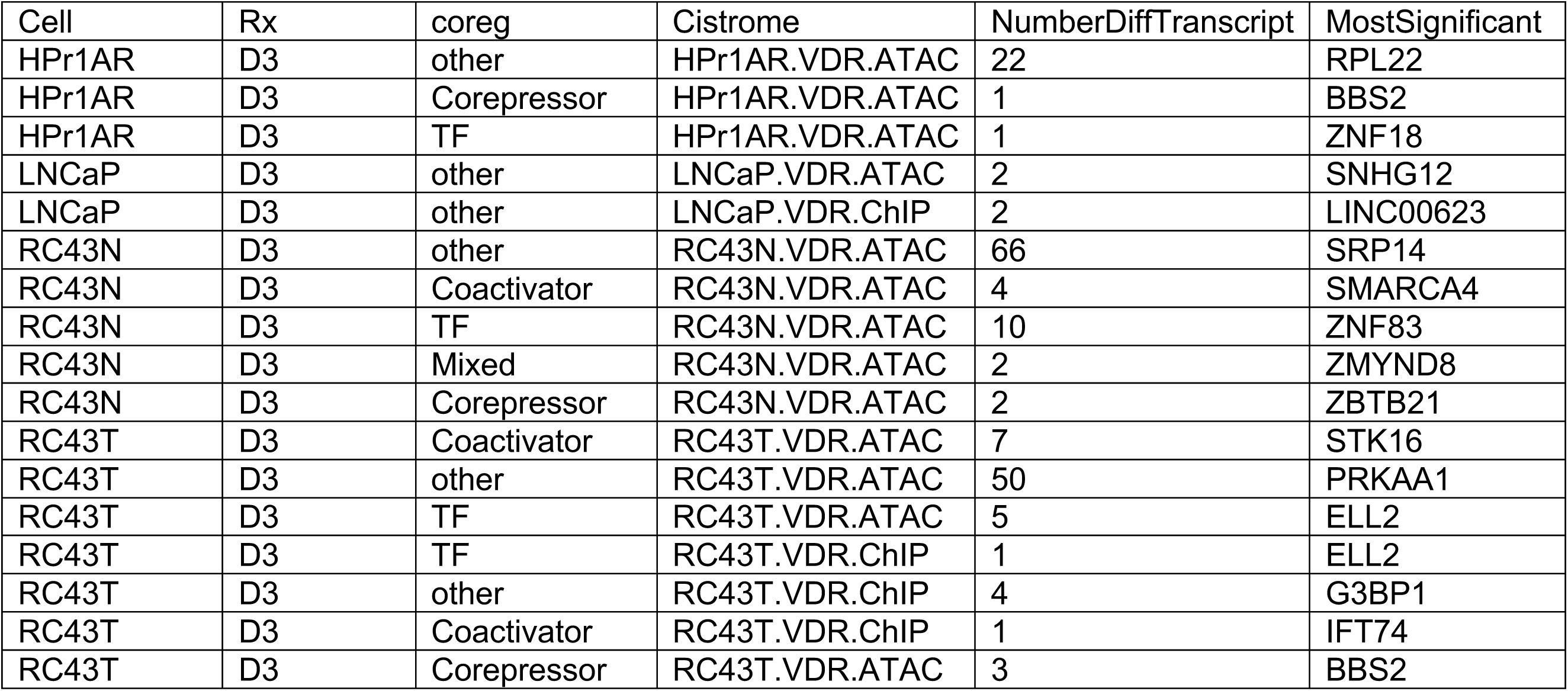
<u>Genes annotated to 1α,25(OH)_2_D_3_-regulated nucleosome free regions and VDR binding sites overlap with differentially expressed transcripts.</u> Genes were annotated to ATAC-Seq or ChIP-Seq regions within 100kB and those genes overlapped with the differentially expressed transcripts, identified by salmon/DRIM-Seq. Overlapped genes were classified either as a Coactivator (CoA), Corepressor (CoR), Mixed function coregulator (Mixed), transcription factor (TF) or other, and the most significant member of each class in each condition is indicated.

Pre-ranked gene set enrichment analysis (GSEA) was undertaken using the Hallmarks, Chemical Perturbations and GO biological terms and underscored how divergent the 1α,25(OH)_2_D_3_- dependent gene expression patterns were between cell types (**Supplementary Figure 8**). HPr1AR and LNCaP cells displayed few significant enrichments (FDR < 0.1), and the only enriched term in LNCaP cells was enrichment of the vitamin D receptor pathway. By contrast, RC43N cells displayed many positively enriched terms including Early Estrogen Response and negatively enriched terms including those associated with cell cycle control. Other modestly positively and negatively enriched gene sets were associated with inflammatory-response, AR and ERα signaling (positive), MYC signaling (negative), and several Circadian rhythm terms. RC43T displayed some similarities (e.g., Estrogen Response signaling and vitamin D pathway), but frequently the terms were either less significant, or enrichment was switched to a negative enrichment.

To test the possibility that indeed GSEA terms (e.g., **Supplementary Figure 9A**) were oppositely enriched, we calculated the delta between NES terms from RC43N and RC43T. Of the 211 terms that were significantly enriched in both RC43N and RC43T, 135 (64.0%) were oppositely regulated with a delta value of greater than 3 (**Figure 4C**) For example, Zhou Cell cycle genes enrichment score (NES) was -2.99 in RC43N, which switched to 1.28 in RC43T. Similarly, terms associated with estrogen signaling, cancer progression, MYC and SMARCE1-dependent targets were oppositely regulated. These findings were complemented by epigenetic landscape in silico deletion analysis, which measures enrichment of transcription factor targeting in the respective transcriptomes[55]. **Supplementary Figure 9B** displays a Euler diagram of the overlap of these transcription factors and demonstrates that although RC43N and RC43T contained distinct set members, but were also essentially overlapping with HPr1AR and LNCaP, supporting the concept that the response in RC43N and RC43T cells is most diverse.

Together these findings reveal that 1α,25(OH)_2_D_3_ transcriptional signaling is more impactful in AA compared to EA prostate models, and that within AA cells, there is frequent gain and loss of enrichment of many pathways suggesting that the capacity of 1α,25(OH)_2_D_3_ signaling is altered.

### Integration of VDR dependent cistromes and transcriptomes reveals stronger gene regulation responses in RC43N and RC43T cells

In the first instance we examined how the cistrome data overlapped by comparing the ATAC-Seq and ChIP-Seq data for each cell and treatment combination. We measured the intersection of the basal and 1*α*,25(OH)_2_D_3_-stimulated VDR ChIP-Seq with the 1*α*,25(OH)_2_D_3_-stimulated mono and NF ATAC-Seq data. Remarkably, none of the cells displayed any significant overlap (a minimum of 1 bp) between ChIP- and 1*α*,25(OH)_2_D_3_-stimulated ATAC-Seq data. That is, although within each cell and condition there were individual genes annotated to have both a significant VDR binding site and a significant 1*α*,25(OH)_2_D_3_-stimulated NF region, these sites did not overlap. By contrast, in RC43N the basal and 1*α*,25(OH)_2_D_3_-stimulated VDR ChIP-Seq data significantly overlapped with the separate basal and 1α,25(OH)_2_D_3_ treated NF ATAC-Seq (i.e., not the differentially enriched 1α,25(OH)_2_D_3_-dependent NF regions) and the basal conditions only in LNCaP (**Supplementary Table 3**) supporting a role for VDR to bind to NF regions in these cells.

Next, each cistrome set was integrated with the transcriptome to identify relationships between either VDR binding or 1α,25(OH)_2_D_3_-induced NF regions, and gene expression, namely peak:gene relationships. To refine these analyses further, the VDR and cistromes were annotated to ChromHMM-defined chromatin states, and annotated to within a 100 kb of the nearest gene, including those that were members of the VDR-interactome as defined by BioGRID [56] (**Supplementary Table 4, 5**). Given there are more 1α,25(OH)_2_D_3_-dependent NF regions, the ATAC-Seq data was annotated to more genes. Within these genes it is interesting to note that RC43N NF regions in ChromHMM defined promoters annotate to the highest number of genes, followed by RC43T. It is also clear that many VDR.biogrid genes are associated with gain NF regions including *TP53* and *NCOR1* in RC43N. Similarly, in RC43T, the loss of NF regions annotates to many VDR BioGRID genes including *LCOR* and *RB*.

By contrast, the basal and 1α,25(OH)_2_D_3_-dependent VDR ChIP-Seq data were annotated to fewer genes, and largest number is in RC43N, where they are evenly distributed evenly between the different ChromHMM regions. Again, reflecting how RC43T cells may lose VDR-genome interactions in the presence of 1α,25(OH)_2_D_3_, almost all peak:gene relationships are in the basal state and most are not annotated to ChromHMM regions. Finally, across the ChIP-Seq data, very few VDR.biogrid genes are identified with notably including LCOR.

We further reasoned that the VDR cistromes represent the sum potential for 1α,25(OH)_2_D_3_- dependent gene expression and therefore restricted VDR-dependent gene:peak relationships to genes detected by regular and small RNA-Seq expressed in the comparisons across AA to EA models (e.g. **Supplementary Figure 7**; AA.N.D3 (RC43N/1α,25(OH)_2_D_3_ compared to HPr1AR/1α,25(OH)_2_D_3_); AA.T.D3 (RC43T/1α,25(OH)_2_D_3_ compared LNCaP/1α,25(OH)_2_D_3_)). Restricting to these genes we then categorized peak:gene relationships for the ChromHMM-classified VDR cistromes and ChromHMM-classified ATAC NF cistromes (**Supplementary Figure 10A**). Next, we further categorized these peak:gene relationships by calculating frequency in 10kb bins up and downstream from DEGs (ChIP-Seq data; **Supplementary Figure 10B**, ATAC-Seq data; data not shown). RC43N gene:peak relationships were most frequent of the models being detected in all classifications in both the basal and 1α,25(OH)_2_D_3_-stimulated states. By contrast, in RC43T the gene:peak relationships were only detected in the basal state. That is, the 30 1α,25(OH)_2_D_3_-stimulated VDR binding sites in RC43T (**Supplementary Table 2**), annotated to 19 genes (**Supplementary Table 4**) but none of these were DEGs identified between RC43T/1α,25(OH)_2_D_3_ compared to LNCaP/1α,25(OH)_2_D_3_.

To test the significance of the VDR-dependent cistrome-transcriptome relationships we applied these classifications and built on the BETA method[57]. Specifically, the sum was calculated of VDR peak or NF scores within 10 kb bins around DEGs multiplied by the absolute gene-specific fold change for each DEG, and weighted to more proximal relationships; this approach was used to test how the summed VDR binding in defined genomic features significantly related to changes in gene expression.

In the first instance, for the ChIP-Seq data (categorized as EtOH-dependent, D3-dependent or shared EtOH.D3-dependent) were integrated with RNA-Seq, regardless of ChromHMM association. This revealed that the few genes in HPr1AR that were bound by 1α,25(OH)_2_D_3_-stimulated VDR were only identified in the positively regulated genes. All other comparisons were compared to RC43N cells, and revealed that in the basal peaks the scores for positively regulated genes in RC43T were significantly greater. This may suggest that basal VDR-bound genes in RC43N are less transcriptionally responsive than the same condition in RC43T. These analyses underscored that lack of a 1α,25(OH)_2_D_3_ VDR-mediated response in RC43T (**Figure 5A**). This was also true considering only genes bound in the Transcribed regions (data not shown).

**Figure 5.**
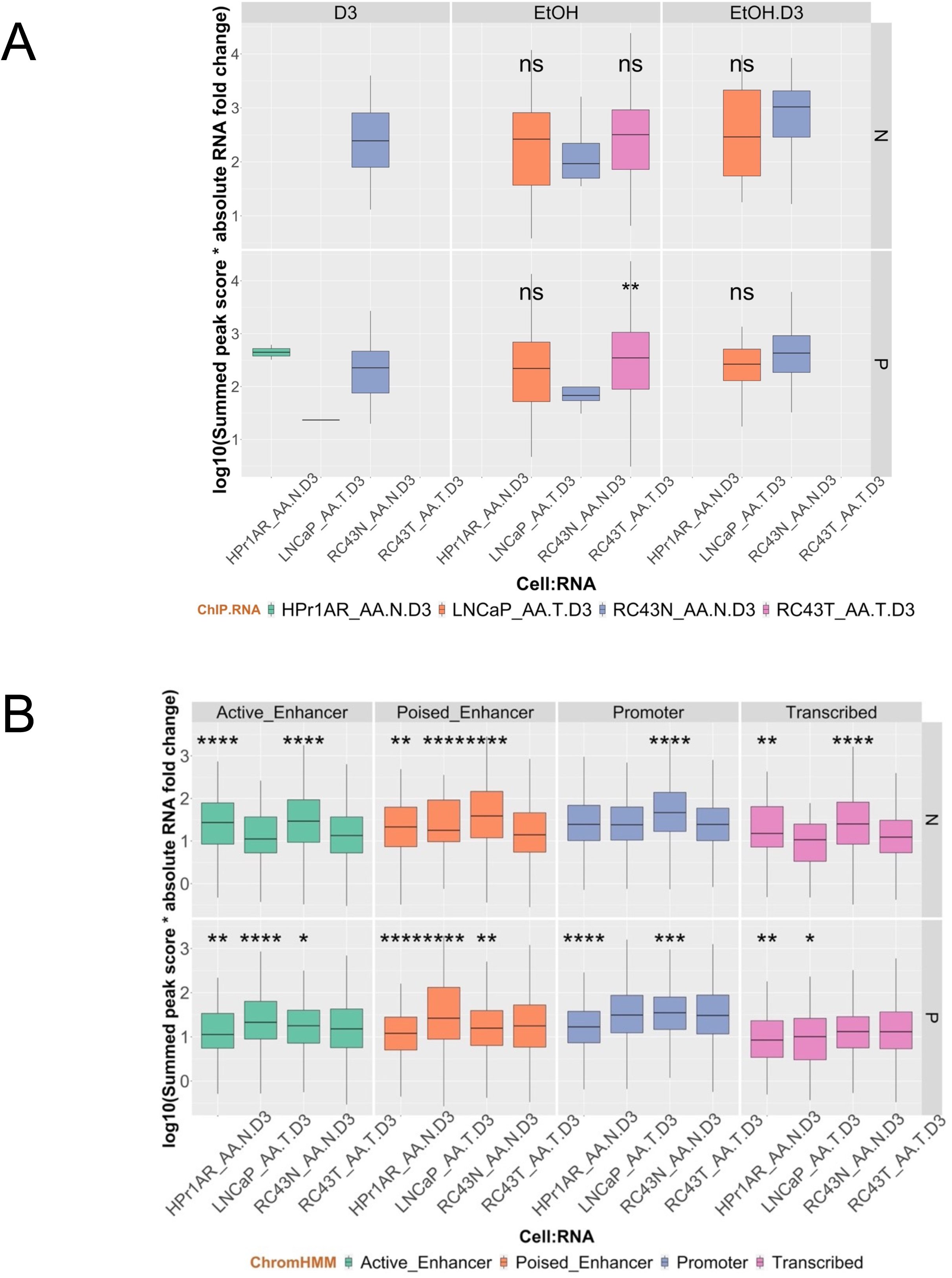
VDR cistrome-transcriptome relationships. **A**. VDR binding sites from the ChIP-Seq data were annotated to DEGs within 100 kb and the sum of peak scores combined with gene expression magnitude were compared across cells, and classified according to ChromHMM distributions. **B**. As above but for ATAC-Seq data.

With more regions in the ATAC-Seq data, and only initiated by 1α,25(OH)_2_D_3_, these relationships were considered in ChromHMM regions. VDR binding and impact across the cells was compared to RC43T and in several cases RC43N summed peak scores were significantly greater, for example Promoters and Poised and Active enhancers, and for negatively regulated genes in Transcribed regions (**Figure 5B**). Together these data support the concept that that VDR binding and 1α,25(OH)_2_D_3_ associated NF regions in RC43N were most consistently associated with stronger patterns of gene expression than the other three cells.

### The BAZ1A/SMARCA5 complex regulates 1α,25(OH)_2_D_3_-stimulated VDR responses

The cistrome-transcriptome studies support the concept that non-malignant RC43N prostate cells are significantly more sensitive to VDR-mediated gene regulation than either RC43T or the EA cells. The VDR binding in RC43N is co-incident with motifs for other nuclear receptors and other factors and including transcription factors that control circadian rhythm. By contrast to RC43N, the isogenic malignant counterpart RC43T, displays suppressed gene expression patterns that include disruption of numerous transcriptional programs. Therefore, we exploited clinical cohorts to determine whether there was evidence for suppressed VDR signaling in AA PCa progression.

In the first instance, we screened a large panel of coregulators[58] in the DEGs between TMPRSS2 fusion positive and negative PCa in EA and AA patients in the TCGA-PRAD cohort. This identified 27 altered coregulators in AA patients, and from these, five were uniquely or more significantly altered in AA compared to EA TMPRSS2 fusion negative PCa (**Figure 6A**). The most altered coregulators included several known to interact with VDR signaling including *BAZ1A* (Bromodomain Adjacent to Zinc Finger Domain 1A), and *SMARCA5/WSTF* (SWI/SNF-Related, Matrix-associated, Actin-dependent Regulator of Chromatin, Subfamily A, Member 5), which functions cooperatively with *BAZ1A*[59]. By contrast, expression of the AR and VDR were unchanged between EA and AA PCa patients. Across the cell models, the expression and regulation of these proteins diverged between EA and AA cell models. BAZ1A expression was highest in LNCaP and unchanging in expression by 1α,25(OH)_2_D_3_, whereas it was strongly induced in RC43N, but strongly repressed in RC43T. These patterns were broadly the same for SMARCA5 (**Figure 6B**).

**Figure 6.**
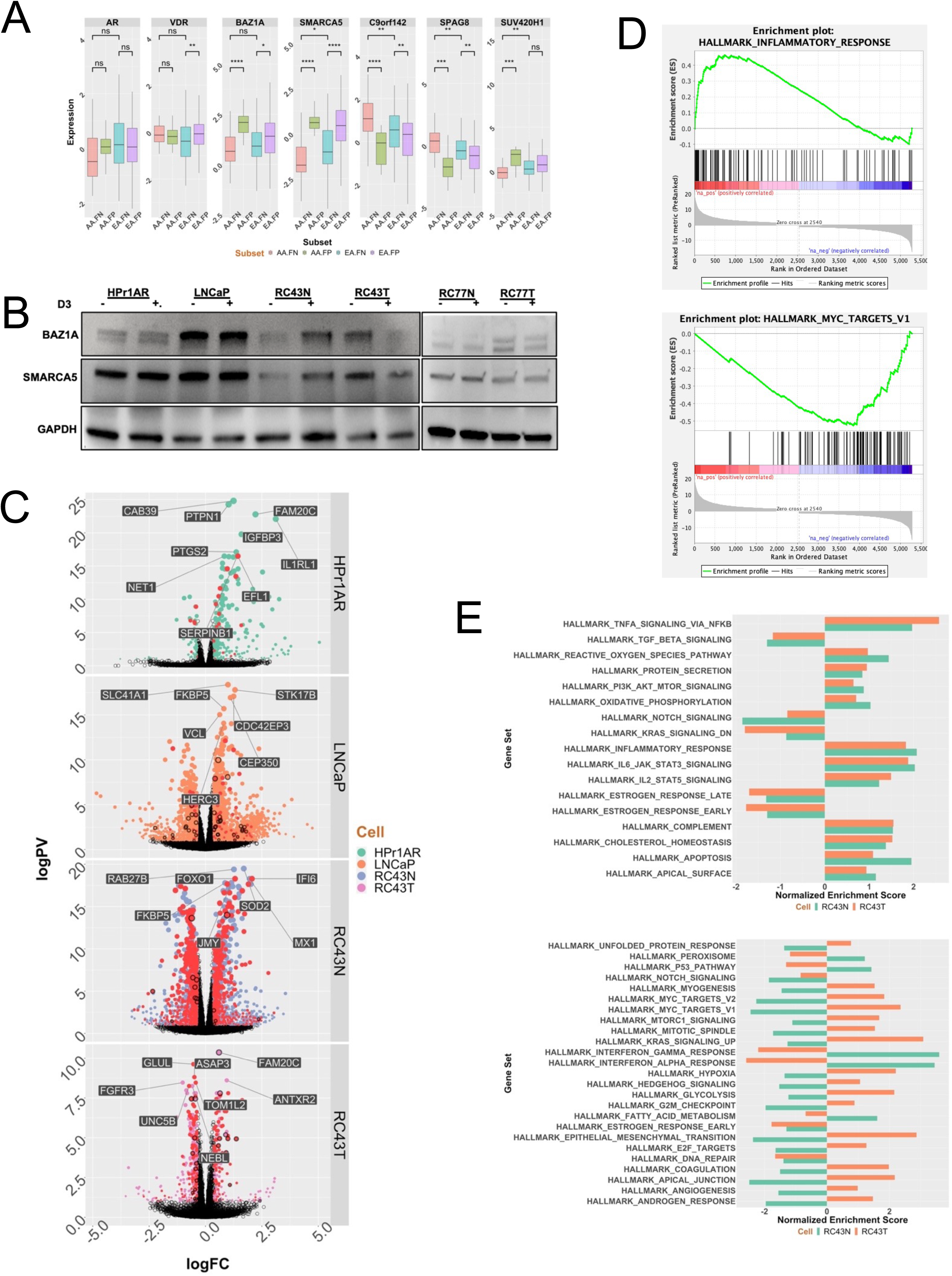
The impact of BAZ1A on expression of VDR-dependent gene networks. **A**. Altered *BAZ1A* and *SMARCA5* expression in African American PCa was identified in the TCGA prostate cancer cohort by comparing EA and AA tumors, and considering status of TMPRSS2 translocations. **B**. Western immunoblot measurements of BAZ1A and SMARCA5 levels after 1α,25(OH)_2_D_3_ treatment (100 nM, 24h) or vehicle control in cell lines. **C**. Unique DEGs in cells with restored expression of BAZ1A. **D**. BAZ1A-dependent DEGs enhanced inflammatory responses and repressed MYC networks. **E**. Comparable and divergent GSEA enrichment in BAZ1A-dependent DEGS in RC43N and RC43T

To identify the impact of altered BAZ1A/SMARCA5 on expression of VDR target genes, we examined the correlations between either BAZ1A or SMARCA5 and the 1α,25(OH)_2_D_3_-regulated genes from RC43N and RC43T in the AA TMPRSS2 fusion negative tumors from the TCGA-PRAD cohort. From these correlations we filtered those genes in Hallmarks_InflammatoryResponse and GO_Circadian Rhythm (**Supplementary Figure 11A**). Supportively, BAZ1A and SMARC5 correlations to these pathway genes were more pronounced for RC43N 1α,25(OH)_2_D_3_-regulated genes than those from RC43T, suggesting that BAZ1A/SMARCA5 is a VDR co-activator and impacts expression of genes associated with inflammation. Next, we examined the genes in BAZ1A containing SWI/SNF complexes that were expressed in the cell lines and tumor cohorts. Expression patterns of the BAZ1A SWI/SNF complex genes significantly distinguished the AA from the EA cell line models (**Supplementary Figure 11B**). The most altered genes in this complex also significantly distinguished *TMPRSS2* fusion positive from negative tumors in the TCGA-PRAD cohort (chi-squared p = 0.002), suggesting that genomic ancestry impacted expression of these genes was most common in the absence of *TMPRSS2* fusion (**Supplementary Figure 11C**). Finally, we examined expression of all the genes in all four SWI/SNF complexes in the Rayford et al cohort of AA tumors (n = 596) compared to EA tumors (n = 556)[10]. All genes detected genes from each of the four complexes were significantly down-regulated in AA tumors compared to EA counterparts (**Supplementary Figure 11D**).

We therefore tested the impact of BAZ1A expression in the four cell line models using RNA-Seq (**Figure 6C, Supplementary Figure 8**). A BAZ1A-containing plasmid was transiently expressed in all four cells lines and elevated expression confirmed (**Supplementary Figure 12A**). In each model we undertook RNA-Seq after 1α,25(OH)_2_D_3_-stimulation (100 nM, 8h) in BAZ1A over-expressed cells compared to vector controls. BAZ1A over-expressed cells displayed enhanced 1α,25(OH)_2_D_3_ responses which was most pronounced in LNCaP and RC43N in terms of the number of genes with enhanced responses. Genes associated with a VDR ChIP-Seq peak or NF regions in the same cell background are represented as a red symbol with a border on the volcano plots (**Figure 6B**).

The VDR cistrome genes were in many cases significantly more enriched in than predicted by chance (**Table 7**). This supports a role for BAZ1A levels to augment regulation of 1α,25(OH)_2_D_3_- regulated genes. However, it is made more striking by the fact that the same analyses of the parental cells (**Figure 4A**) found no evidence for significant enrichment.

**Table 7:** <u>Overlap of BAZ1A-dependent 1α,25(OH)_2_D_3_-regulated gene expression with genes annotated to 1α,25(OH)_2_D_3_-regulated nucleosome free regions and VDR binding sites significantly.</u> Genes were annotated to ATAC-Seq or ChIP-Seq regions within 100kB and those genes overlapped with the differentially expressed genes in the same cell background was BAZ1A was overexpressed, and enrichment tested with a hypergeometric test (lower.tail = FALSE).

| Cell | Cistrome | logPval | NumberCistromeGenes | Threshold | MostSignificant |
| --- | --- | --- | --- | --- | --- |
| RC43N | RC43N.VDR.ATAC | 11.35 | 1358 | Significant | PIK3R6 |
| RC43T | RC43T.VDR.ATAC | 3.01 | 349 | Significant | PIK3C2G |
| RC43T | RC43T.VDR.ChIP | 1.40 | 28 | Significant | FRG2B |
| HPr1AR | HPr1AR.VDR.ATAC | 0.82 | 73 | NS | KCNH6 |
| LNCaP | LNCaP.VDR.ATAC | 0.40 | 80 | NS | ANGPTL7 |
| RC43N | RC43N.VDR.ChIP | 0.28 | 14 | NS | ANKRD26P1 |
| LNCaP | LNCaP.VDR.ChIP | 0.26 | 51 | NS | POTEM |

Analyses of the GSEA terms also supported a role for BAZ1A to impact gene expression. For example, in LNCaP cells the combined effect of BAZ1A/1α,25(OH)_2_D_3_ elevated enrichment of genes associated with the Androgen response, UV response and decreased enrichment of MYC targets. By contrast, in RC43N the most enriched terms were Interferon *γ* and *α*, Inflammatory responses, IL6 signaling and TNFA signaling suggesting an immunomodulatory phenotype (**Figure 6 D**). Focusing on the impact of BAZ1A on 1α,25(OH)_2_D_3_-induced expression changes in RC43N and RC43T, and calculating the enrichment terms that change the most, in either the same or a divergent manner, revealed that BAZ1A has a potent and cell-specific impact on gene expression patterns induced by 1α,25(OH)_2_D_3_ (**Figure 6 E**). For example, commonly enriched terms included many of the inflammation terms such as IL-6- and IL-2-mediated signaling, and TNFA signaling as well as Estrogen signaling and apoptosis pathways, whereas divergent pathways included interferon signaling, MYC signaling and androgen responses.

### VDR cistrome-transcriptome target genes in AA PCa clinical cohorts

Finally, we examined how these cistrome-transcriptome relationships established AA and EA cell lines could be detected in clinical cohorts of AA and EA men with PCa, using three cohorts. In the first instance, we examined serum samples from patients with high-grade prostatic intraepithelial neoplasia (HGPIN) who participated in a Southwest Oncology Group (SWOG) clinical trial (SWOG S9917) [60]. Within the cohort, 40% of men progressed from HGPIN to PCa and we examined miRNA that significantly predicted HGPIN progression to PCa in either EA or AA enrolled patients. In this manner we identified separate miRNA associated with HGPIN progression to PCa, and the top 20 in AA patients and EA patients are shown **Supplementary Table 6**. Twelve of the 33 miRNA (36%) that associated with AA progression were annotated to AA 1α,25(OH)_2_D_3_/VDR cistrome regions including *MIR23B* and *RTCA* (contains *MIR553*) (**Table 8**), whereas this was 37/280 (13%) for the EA progression miRNA.

**Table 8:** <u>Serum expression of miRNA that associate with progression from high-grade prostatic intraepithelial neoplasia to PCa in AA and EA patients.</u> Serum samples from 96 patients with high-grade prostatic intraepithelial neoplasia (HGPIN) who participated in a Southwest Oncology Group (SWOG) clinical trial (SWOG S9917). The cohort consisted of 21 AA patients (9 progressed to PCa) and 75 EA patients (33 progressed to PCa). Nanostring PCR was used to measure differential miRNA expression within AA or EA samples and between EA and AA samples by progression status. The data were processed with NanoStringDiff and significantly different miRNA identified (logPV > 1 & absFC > .58). 33 miRNAs significantly associated with AA progression to PCa and more than 200 EA miRNA associated with PCa progression. The AA and EA miRNA genes (or host genes) that exclusively associated with progression were annotated to AA or EA 1α,25(OH)_2_D_3_/VDR cistrome regions, and the number of annotations is indicated and an example of the most significantly regulated miRNA given.

| Progression | Cistrome | NumberMIRNA | MostSignificant.MIRNA |
| --- | --- | --- | --- |
| AA_Prog | indep | 21 | MIR19A |
| AA_Prog | RC43T.VDR.ChIP | 1 | MIR23B |
| AA_Prog | RC43N.VDR.ATAC | 4 | MIR553 |
| AA_Prog | RC43T.VDR.ATAC | 7 | MIR553 |
| EA_Prog | indep | 243 | MIR412 |
| EA_Prog | LNCaP.VDR.ChIP | 2 | MIR664A |
| EA_Prog | HPr1AR.VDR.ATAC | 32 | MIR2110 |
| EA_Prog | LNCaP.VDR.ATAC | 3 | MIR3605 |

Next, we re-examined data from our earlier PCa RNA-Seq study from EA and AA patients who received vitamin D_3_ (4000 IU daily) prior to radical prostatectomy. Ancestry informative markers confirmed the African genomic ancestry of the AA patients (data not shown). As we reported previously [7] the responses in EA patients were essentially null, whereas a strong transcriptional response was observed in the AA patients with ∼1400 genes being significantly regulated by vitamin D_3_, and GSEA revealed these genes were enriched in immuno-modulatory and PCa-relevant pathways (**Figure 7 A, B**). Remarkably, 326 (∼23%) of these genes were significantly bound by VDR in ChIP-Seq studies. The DEGs were again enriched for inflammatory signaling components. Interestingly, 70% of the significantly modulated genes in the AA patients were either VDR ChIP-Seq or ATAC-Seq annotated genes (**Table 9**)

**Figure 7.**
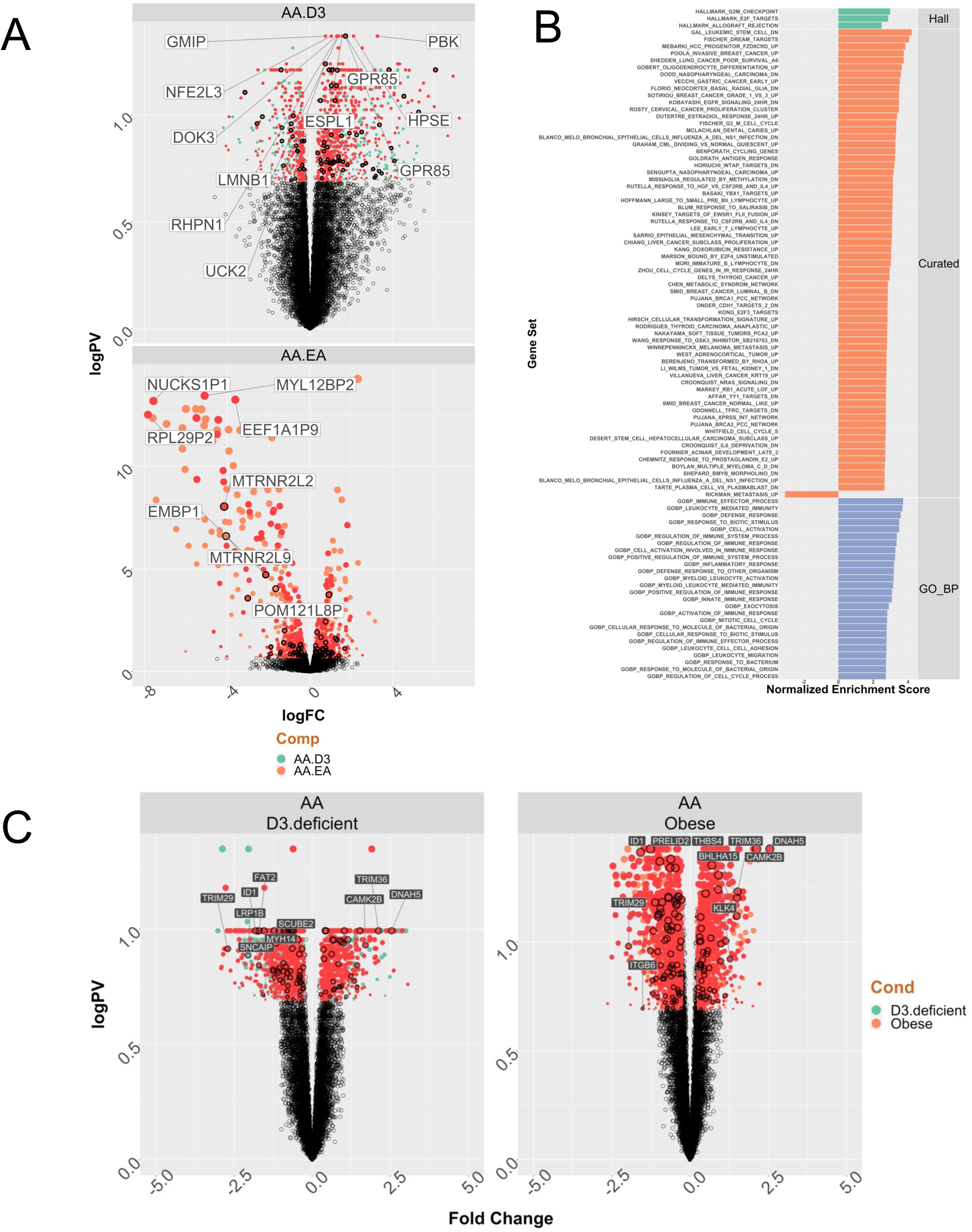
VDR cistrome-transciptome relationships in prostate cancer. **A**. A cohort of seven AA PCa patients with conserved African genomic ancestry and 16 EA PCa patients were treated with vitamin D_3_ (4000 IU daily), and RNA-Seq undertaken on the tumors following radical prostatectomy, as we reported previously[7]. Significantly differentially regulated genes in the AA PCa group (there were no DEGs in the EA PCa group) were overlapped with genes annotated to ATAC-Seq or ChIP-Seq regions within 100kB. The Volcano plot of the DEGs for the response in AA men, or comparing basal AA to EA PCa and annotated with genes that are VDR bound and/or 1α,25(OH)_2_D_3_-dependent NF region annotated genes. **B**. GSEA analyses of the vitamin D_3_ stimulated AA tumors **C**. RNA-Seq was undertaken in tumors from a cohort of 57 AA and 18 EA patients who underwent radical prostatectomy. Tumor-specific significant DEGs were identified for deficient serum 25(OH)D_3_ (serum 25(OH)D_3_ levels < 12 ng/ml; low) or obesity (BMI > 30; O). In each case BMI or 25(OH)D_3_ levels were kept as a continuous variable respectively. The DEGs for 25(OH)D_3_ deficiency (left) of obesity (right) in the AA PCa group (there were no DEGs in the EA PCa group) were overlapped with genes annotated to ATAC- Seq or ChIP-Seq regions within 100kB.

**Table 9:** <u>Percentage overlap of vitamin D_3_-regulated gene expression in prostate tumors from AA patients with genes annotated by 1α,25(OH)_2_D_3_-regulated nucleosome free regions and/or VDR ChIP-Seq.</u> A cohort of seven AA PCa patients with conserved African genomic ancestry and 16 EA PCa patients were treated with vitamin D_3_ (4000 IU daily), and RNA-Seq undertaken on the tumors following radical prostatectomy, as we reported previously[7]. Significantly differentially regulated genes in the AA PCa group (there were no DEGs in the EA PCa group) were overlapped with genes annotated to ATAC-Seq or ChIP-Seq regions within 100kB. The percentage overlap of DEGs and cistrome genes is shown.

| AA.ChIP.targets | AA.ATAC.targets | NumberTargets | Percent |
| --- | --- | --- | --- |
| AA.ChIP | AA.ATAC | 55 | 70.5 |
| AA.ChIP | indep | 23 | 29.5 |
| indep | AA.ATAC | 950 | 69.2 |
| indep | indep | 423 | 30.8 |

Finally, in a further cohort of 57 AA and 18 EA patients we undertook RNA-Seq in tumor and contralateral normal material collected from clinics in Chicago, IL [61]. DEG analyses were undertaken in tumors and normal material in two comparisons to identify the race-specific impact of serum 25(OH)D_3_ levels and obesity. Firstly, we compared how gene expression differed between in the cohort with deficient serum 25(OH)D_3_ levels (< 12 ng/ml) compared to replete levels, while keeping obesity as a continuous variable. Secondly, we compared how gene expression differed between the obese and normal groups, while keeping serum 25(OH)D_3_ levels as a continuous variable.

In the AA cohort there were 1612 DEGs in the tumors with low serum 25(OH)D_3_ levels compared to tumors from patients with replete levels. Reflecting, the radical prostatectomy samples, there were no significant DEGs in EA patients when considering the levels of serum 25(OH)D_3_. Similarly, in the AA cohort comparing gene expression in tumors from men who are obese compared to normal counterparts there were 2415 DEGs; again, there were no DEGs in the EA tumor group between obese and normal BMI patients. Finally, these DEGs were queried as to whether they were also annotated to be VDR ChIP-Seq or 1α,25(OH)_2_D_3_ stimulated NF ATAC-Seq genes (**Figure 7C**). **Table 10** summarizes the overlaps of these DEGs and cistrome data with 85% of either DEG group (impact of serum 25(OH)D_3_ or obesity) overlapping with the AA cistrome data.

**Table 10:**
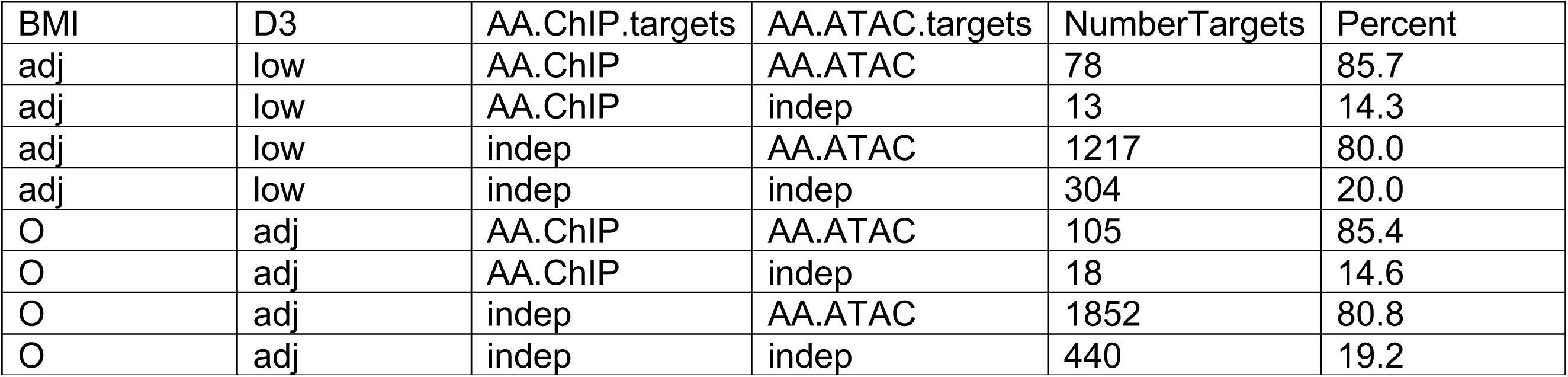
<u>Percentage overlap of either serum 25(OH)D_3_ vitamin D_3_-regulated or obesity-regulated gene expression in prostate tumors from AA patients with genes annotated by 1α,25(OH)_2_D_3_-regulated nucleosome free regions and/or VDR ChIP-Seq.</u> RNA-Seq was undertaken in tumors from a cohort of 57 AA and 18 EA patients who underwent radical prostatectomy. Tumor-specific significant differentially expressed genes (DEGs) were identified for deficient serum 25(OH)D_3_ (serum 25(OH)D_3_ levels < 12 ng/ml; low) or obesity (BMI > 30; O). In each case BMI or 25(OH)D_3_ levels were kept as a continuous variable respectively (adj). The DEGs in the AA PCa group (there were no DEGs in the EA PCa group) were overlapped with genes annotated to ATAC-Seq or ChIP-Seq regions within 100kB. The percentage overlap of DEGs and cistrome genes is shown.

| BMI | D3 | AA.ChIP.targets | AA.ATAC.targets | NumberTargets | Percent |
| --- | --- | --- | --- | --- | --- |
| adj | low | AA.ChIP | AA.ATAC | 78 | 85.7 |
| adj | low | AA.ChIP | indep | 13 | 14.3 |
| adj | low | indep | AA.ATAC | 1217 | 80.0 |
| adj | low | indep | indep | 304 | 20.0 |
| O | adj | AA.ChIP | AA.ATAC | 105 | 85.4 |
| O | adj | AA.ChIP | indep | 18 | 14.6 |
| O | adj | indep | AA.ATAC | 1852 | 80.8 |
| O | adj | indep | indep | 440 | 19.2 |

## DISCUSSION

The current study aimed to define if 1α,25(OH)_2_D_3_ and VDR signaling capacity could provide insights about biological mechanisms involved in racial disparities in PCa between African American and European American menStrong epidemiological findings, as well as clinical trial data broadly support the concept that VDR signaling activated by 1α,25(OH)_2_D_3_ exerts a tumor suppressive effect in AA PCa patients, and potentially that this patient group is also most acutely vulnerable to a lack of 1α,25(OH)_2_D_3_ activated VDR signaling. However, genomic insight into these relationships is not well established, and our proteogenomic study sought to fill this knowledge gap.

To define VDR actions across AA and EA prostate models we generated and integrated high dimensional genomic, transcriptomic and proteomic data sets, combined with clinical genomic data to allow a comprehensive insight into how VDR signaling is altered within and across races. Our findings support several key insights and inferences on VDR signaling, and perhaps are also relevant to a larger set of nuclear receptors in the context of health disparities. Furthermore, to the best of our knowledge, this study represents the first comparative cistromic study between AA and EA cell models and these findings may be insightful to other workers addressing cancer and race-related health disparities.

A starting point for our study was to consider expression of the VDR protein and capture its interactome, which established that while prominent VDR expression is shared in the EA and AA cell models, the interacting proteins and cellular 1α,25(OH)_2_D_3_ sensitivity varies widely. For example, the VDR complex differed by how it was associated with coactivators such as SMARCA5, splicing factors such as DDX39B, and the transcriptional regulator TRIM29. Beyond this, proteins that are also part of the regulation of circadian rhythm were enriched in the VDR complex, such as a gain of ARNTL2 and loss of NONO in RC43T cells, supporting a role for VDR to cooperate with the control of circadian rhythms.

Next, we considered the genomic impact of 1α,25(OH)_2_D_3_ by examining the overlap in nucleosome positioning and where the VDR binds. These approaches established that 1α,25(OH)_2_D_3_’s genomic impact is greatest in RC43N and RC43T, but divergent between these cells. 1α,25(OH)_2_D_3_- stimulated NF regions in RC43N occurred more frequently than in other models, were largely unique and >98% represented a gain. In contrast, 1α,25(OH)_2_D_3_-stimulated NF regions in RC43T were less frequent and resulted in an overall NF region loss. This suggests that VDR is primed to produce very different responses between these two cells, although motif enrichment suggested that the TFs that are enriched in these regions are similar and include AP-1 factors, such as KLF5. The divergence of this response was illustrated by changes in enrichment of circadian rhythm TF motifs, such as CLOCK.

Similarly, ChIP-Seq revealed strong agreement for where the VDR bound in the absence of 1α,25(OH)_2_D_3_ across LNCaP, RC43N and RC43T with more than 300 shared binding sites. The addition of 1α,25(OH)_2_D_3_ resulted in distinct patterns of VDR binding within a cell, and reduced the number of shared sites with only nine sites in common across all cells. Furthermore, reflecting the NF data, there was a dramatic reduction in VDR binding in RC43T. Motif enrichment supported the concept that the basal VDR state in LNCaP and RC43T was the most TF active state with a clear loss of motif enrichment when 1α,25(OH)_2_D_3_ was present. In RC43T, the loss of VDR binding reflected the loss of NF regions and again suggests that the system is altered to limit the impact of 1α,25(OH)_2_D_3_. Reflecting the genomic data, at the transcriptional level the RC43N and RC43T cells were most responsive and indeed many genes that were modulated by 1α,25(OH)_2_D_3_ also were either bound by VDR, or had a change in 1α,25(OH)_2_D_3_-stimulated NF distribution. Again, the response between RC43N and RC43T were frequently opposed, with many GSEA terms being switched in direction between the cells.

Together these data support the concept that RC43N and RC43T have the strongest response to 1α,25(OH)_2_D_3_ but the effect is highly divergent between them. We therefore sought mechanisms that may drive this divergence. Analyses of TCGA data identified a potential role for altered expression of the BAZ1A/SMARCA5 SWI/SNF complex. Therefore, we tested this possibility. Again, underscoring the differences between RC43N and RC43T, 1α,25(OH)_2_D_3_-treatment upregulated BAZ1A and SMARCA5 in RC43N, but down-regulated these proteins in RC43T. It is noteworthy that SMARCA5 was significantly less enriched in the VDR complex in RC43T compared to LNCaP, that RC43N VDR ChIP-Seq significantly overlapped with publicly available SMARCA4 ChIP-Seq from LNCaP cells, and that 1α,25(OH)_2_D_3_ treatment resulted in isoform-specific expression of SMARCA4 in RC43N. It is also worth noting that GWAS SNPs in BAZ1A associate significantly with heel bone strength, also supporting a role in regulating VDR function [62]. Restoring BAZ1A expression led to significantly enhanced 1α,25(OH)_2_D_3_-regulated transcriptome, but these genes were only significantly enriched for VDR bound and NF regions in RC43N and RC43T suggesting that BAZ1A expression had the most significant impact on VDR function in AA cells. The most strongly up-regulated gene in RC43N that is annotated to RC43N 1α,25(OH)_2_D_3_-regulated NF region was *PIK3R6*, which recently was identified as a novel antigen in a clinical immunotherapy trial in advanced PCa[63]. These data also contribute to the earlier analyses of VDR functions in the prostate[64] and also mechanisms by which VDR limits proliferation[65].

By identifying AA and EA specific VDR genomic events, we measured if these differences were apparent in three clinical cohorts. These analyses revealed that the footprint of VDR signaling was most apparent in AA PCa. For example, miRNA that predicted progression from HGPIN to PCa in AA men were highly enriched for VDR cistrome data, as were genes that responded in prostate tumors from men receiving vitamin D3 supplementation prior to radical prostatectomy. This was also strikingly apparent in prostate tumors from men who had deficient serum 25(OH)D_3_ levels, and indeed in this interacted significantly with obesity (BMI >30.0 kg/m^2^) status.

It has been clear for many years [66] that the vitamin D signaling axis has systemic functions beyond the control of calcium regulation. Indeed, large scale transcriptomic and proteomic studies have revealed an interesting paradox, namely that the VDR is expressed widely and robustly across cell-types including those that have no obvious role in calcium function [67, 68]. Furthermore the impact of altered vitamin D levels is strongly implicated in the etiology of non-calcemic syndromes such as diabetes cardiovascular disease, autoimmune disease, and various high-profile cancers. Recently, in the VITAL cohort, vitamin D3 has been shown to reduce the incidence of autoimmune diseases [69].

Although guidance is available for serum 25(OH)D_3_ levels required for bone health, it is far from clear what level is required either to promote cardiovascular health or to prevent autoimmune diseases. It is even less clear how vitamin D deficiency amongst African Americans, which is highly prevalent, impacts cancer and these other diseases. Our genomic data suggest that the powerful example of changing melanin content in the skin through adaptation has occurred in parallel with a range of cellular mechanisms that include the genomic functions of the VDR in non-calcemic tissues.

Consistently, through this study, the data support a model whereby VDR signaling is more potent in AA prostate cell models and prostate tumors. VDR-induced enrichment of circadian rhythm and inflammatory networks are evident. Observational evidence supports a cross-talk between VDR and components of the transcriptional network that regulates melatonin production, circadian rhythm and sleep duration[40, 70]. Given the manner in which VDR function can be corrupted, either through changes in serum Vitamin D3 levels[15, 16, 30, 71, 72], changes in membrane transport[27] or disruption to the composition of the VDR complex [73–81], it is reasonable to infer that the VDR stands at the crossroads of biopsychosocial signaling that impacts PCa by a distinct pattern of VDR genomic binding in cells with African genomic ancestry. That these VDR patterns in prostate cells are distinctly enriched for genes that control of circadian rhythm and inflammatory networks suggest that men of African ancestry may be disproportionally impacted by environments of low UVB exposure, increase stress, access to quality early detection and treatment, as well work environments that are deleterious for circadian signaling. To this concept we have added the mechanistic insight that the SWI/SNF complexes containing BAZ1A and SMARCA5 are distorted in a manner that reflects genomic ancestry, and further distorts the normal functions of the VDR and suggests enhanced sensitivity to DNA damage[82].

In conclusion, the current study suggests that the functions of the VDR may have adapted with significant distinctions between people of different genomic ancestry. More specifically, we reason that adaptation to environments of lower UVB exposure that could potentially be associated with vitamin D insufficiency and alter the prominence of how VDR signaling occurs in a wide variety of tissues. Furthermore, we propose that the prostate is an important gland with which to test this possibility given it is the site of significant syndromes and diseases that are highly impactful on US men, and furthermore many of these conditions display significant health disparities.

## MATERIALS AND METHODS

### Cell culture and materials

The EA cells HPr1AR, LNCaP (*140*) and the AA cells RC43N and RC43T were maintained at 37°C and 5.0% CO2 using a cell culture incubator with UV contamination control (Sanyo). HPr1AR cells were maintained in keratinocyte serum free media (supplemented with 25mg of Bovine pituitary extract, 10ug epithelial growth factor and 10% FBS); LNCaP cells in RPMI 1640 Medium containing 10% FBS; RC42N and RC43T in keratinocyte serum free media (supplemented with 25mg of Bovine pituitary extract, 10ug epithelial growth factor and 10% FBS). All media was supplemented with 100 U/mL Penicillin-Streptomycin. 1α,25(OH)_2_D_3_ was kept as 10mM EtOH stocks, and diluted to 1000x stocks prior to treatments. Cell lines were authenticated by STR profiling and confirmed mycoplasma free by RT-PCR.

### RT-qPCR

Total RNA was isolated via TRIzol® reagent (Thermo Fisher Scientific) for candidate mRNA detection by use of the AllPrep DNA/RNA/miRNA Universal Kit (Qiagen), following manufacturer’s protocols. Complementary DNA (cDNA) was prepared using iScriptTM cDNA Synthesis Kit (Bio-Rad), following manufacturer’s protocols. Relative gene expression was subsequently quantified via Applied Biosystems 7300 Real-Time PCR System (Applied Biosystems), for both TaqMan and SYBR Green (Thermo Fisher Scientific) applications. All targets were detected using either pre-designed TaqMan Gene Expression Assays (Thermo Fisher Scientific; *VDR, BAZ1A, SMARCA5* using pre-designed PrimeTime qPCR primers and a final primer concentration of 500nM. All primers for use with SYBR Green application were tested for specificity by melting curve analysis with subsequent product visualization on agarose gel. All RT-qPCR experiments were performed in biological triplicates, with at least technical duplicates. Fold changes were determined using the 2-ΔΔCt method as the difference between experimental group and respective control group. Significance of experimental comparisons was performed using Student’s t-test.

### Stable transfection of BAZ1A

GFP-BAZ1A was purchase from Addgene (plasmid # 65371)[83] and transfection was cells were transfected and stable selection after transduction with by selection and maintainance in media supplemented with puromycin (2µg/mL).

### Western Immunoblotting

Total cellular protein was isolated from exponentially growing cells for determination of target protein expression. Cells were harvested, then washed in ice cold PBS before lysing in ice cold RIPA buffer (50mM Tris-HCl pH 7.4, 150mM NaCl, 1% v/v Triton X-100, 1mM EDTA pH 8.0, 0.5% w/v sodium deoxychlorate, 0.1% w/v SDS) containing 1x cOmplete Mini Protease Inhibitor Tablets (Roche). Protein concentrations were quantified using the DC Protein Assay (Bio-Rad), following manufacturer’s protocols. Equal amounts of proteins (30-60µg) were separated via SDS polyacrylamide gel electrophoresis (SDS-PAGE) using precast 4-20% gradient polyacrylamide gels (Mini-Protean TGX, Bio-Rad). Proteins were transferred onto polyvinylidene fluoride (PVDF) membrane (Roche) for 30V for 16 hours. Post transfer, membranes were blocked with 5% non-fat dry milk (NFDM) for 1 hour at room temperature. Blocked membranes were probed with primary antibody against BAZ1A, SMARCA5, VDR, GAPDH either overnight at 4°C or for 3 hours at room temperature. Primary antibody was detected after probing for 1 hour with HRP-linked rabbit anti-mouse IgG (P0161, Dako) or goat anti-rabbit IgG (P0448, Dako) secondary antibody at room temperature using ECL Western Blotting substrate (Pierce). Signal quantification was performed using the ProteinSimple Fluorochem M Imager.

### Cell viability

Bioluminescent detection of cellular ATP as a measure of cell viability was undertaken using CellTiter-Glo^®^ (promega) reagents. Cells were plated at optimal seeding density to ensure exponential growth in 96-well, white-walled plates. Wells were dosed with agents to a final volume of 100 µl. Dosing occurred at the beginning of the experiment, and cells were incubated for up to 120 hours. Luminescence was detected with Synergy^TM^ 2 multi-mode microplate reader (BioTek® Instruments). Each experiment was performed in at least triplicate wells in triplicate experiments.

### Clonogenic assays

The colony formation ability of cells in presence of variable doses of 1α,25(OH)_2_D_3._ 1000 cells were plated in triplicates in a 6 well plate and treated with 1α,25(OH)_2_D_3_ every three days for a period of 14days. After 14 days cells were washed and fixed with neutral buffered formaling and stained with crystal violet stain and quantified as previously reported[84].

### Rapid immunoprecipitation mass spectrometry of endogenous protein

RIME analyses were undertaken with antibody towards the VDR in cells treated with either vehicle or 1α,25(OH)_2_D_3_. 20x10^6^ cells were crosslinked with 1% formaldehyde solution, quenched with glycine (0.1 M, pH 7.5) and harvested in cold PBS. Nuclei were separated as previously reported [85] and subjected to sonication (30s on 30s off cycles for 30mins) for genomic DNA fragmentation. 30ul of 10% triton-X was added and high-speed centrifugation was performed to separate the nuclear proteins. Further, VDR and IgG antibody conjugated beads were incubated with nuclear lysates overnight and washed ten times with RIPA buffer and 2 washes of Ambic solution as previously described [85]. LC-MS/MS was performed over a 2-hour separation and mean spectral count results were analyzed with a generalized linear model workflow to identify differentially enriched proteins.

### RNA-Seq

RNA was extracted from cells in the presence of 1α,25(OH)_2_D_3_ (100nM, 8hr) or EtOH in biological triplicate samples and analyzed by RNA-Seq. Sequencing was performed at the Nationwide Children’s Hospital Institute for Genomic Medicine, and sequencing libraries prepared with the TruSeq Stranded Total RNA kit (Illumina Inc), from 1ug total RNA. Alignment of raw sequence reads to the human transcriptome (hg38) was performed via Rsubread [86] and transcript abundance estimates were normalized and differentially expressed genes (DEGs) identified using a standard edgeR pipeline. Functional annotation of gene sets: Pathway enrichment analysis and gene set enrichment analysis (GSEA) were performed using gene sets from the Molecular signatures database (MSigDB).

### Small RNA-Seq

Cell lines were treated as RNA-Seq and sequencing performed at Nationwide Children’s Hospital Institute for Genomic Medicine. Sequencing libraries were prepared with the TruSeq Small RNA kit (Illumina Inc), from 1ug total RNA. Following manufacturer’s protocols, ligation of 5’ and 3’ RNA adapters to the mature miRNAs 5ʹ-phosphate and 3ʹ-hydroxyl groups, respectively was undertaken. Following cDNA synthesis, the cDNA was amplified with 11-13 PCR cycles using a universal primer and a primer containing one of 48 index sequences, which allowed pooling of libraries and multiplex sequencing. Prior to pooling, each individual sample’s amplified cDNA construct was visualized on a DNA-HS Bioanalyzer DNA chip (Agilent Technologies) for mature miRNA and other small RNA products (140-150bp). Successful constructs were purified using a Pippen prep (Sage Inc.), using 125 – 160 bp product size settings with separation on a 3% agarose gel. The purified samples were validated for size, purity and concentration using a DNA-HS Bioanalyzer chip. Validated libraries were pooled at equal molar to a final concentration of 10nM in Tris-HCI 10 mM, pH 8.5, before 50 cycle sequencing on a MiSeq (Illumina, Inc.). Fastq files were aligned to the genome (hg38) using Rsubread. Expression counts were called against the miRbase consensus miRnome using featureCounts and A standard edgeR pipeline determined differentially expressed miRNA.

### ChIP-Seq

ChIP was performed cells in the presence of 1α,25(OH)_2_D_3 2_D_3_ (100nM, 6hr) or EtOH in triplicate or EtOH in triplicate independent experiments. Briefly, approximately 20x10^6^ cells were crosslinked with 1% formaldehyde solution, quenched with glycine (0.125 M) and harvested in cold PBS. Sonication of crosslinked chromatin was performed using a Bioruptor® UCD-200^TM^ Sonicator (Diagenode) with optimized cycles for each cell type. Immunoprecipitation of sonicated material was performed with antibodies against VDR (D2K6W) – Cell Signaling) or IgG (Rabbit IgG sc2729 – Santa Cruz) for 16 hours, and antibody/bead complexes isolated with Magna ChIPTM Protein A+G magnetic beads (Millipore). Complexes were washed, reverse crosslinked, and treated sequentially with RNase and proteinase K prior to DNA isolation. Sequencing was performed at the Nationwide Children’s Hospital Institute for Genomic Medicine. The VDR cistrome was analyzed with Rsubread/csaw [87] along with TF motif analyses (MotifDb).

In order to find potential transcription factor binding enrichment within VDR cistromes, GIGGLE was utilized to query the complete human transcription factor ChIP-seq dataset collection (10,361 and 10,031 datasets across 1,111 transcription factors and 75 histone marks, respectively) in Cistrome DB[51]. Prostate specific filtering limited analysis to 681 datasets across 74 TFs and 238 datasets across 19 HMs. For each query dataset, we determined the overlap of each VDR cistrome. Putative co-enriched factors were identified by assessment of the number of time a given factor was observed in the top 200 most enriched datasets relative to the total number of datasets for that factor in the complete Cistrome DB (> 1.2 FC enrichment over background). For prostate specific analysis, overlaps across datasets were averaged for each factor.

### ATAC-Seq

50000 cells were plated and treated with 1α,25(OH)_2_D_3_ and vehicle control for 4h and cells were washed twice with PBS and collected by trypsinization. Cells were resuspended in 50ul of ATAC-resuspension buffer (ATAC-RSB - 10mM Tris-HCl, 10mM NaCl, 3mM MgCl_2_) containing (0.1% NP-40, 0.1% tween-20, and 0.01% digitonin) and pipetted up and down 3 times. Further, 1ml of ATAC-wash-resuspension buffer (ATAC-RSB + 0.1% tween 20) was used to pellet down the nuclei. The nucleic were further resuspended in transposition mix (2X TD buffer, 1X PBS, Digitonin 0.01%, tween 20 0.1%, NFW 5ul, and Illumina transposase 2.5ul). Mixing, cleanup and library preparation, quantification and sequencing was performed using NovaSeq6000 S1 PE150bp Sequencing as per protocol (28846090). (ATAC-Seq data were separated into nucleosome free (NF), mono-, di- and tri-nucleosome compartments (ATACSeqQC)[88].

### MiRNA expression in serum samples

In serum samples from SWOG S9917 (high-grade prostatic intraepithelial neoplasia (HGPIN) to PCa), Nanostring was used to identify miRNA associated with progression in AA patients. Nanostring PCR was used to measure differential miRNA expression within race and across race by progression status. The data were processed with NanoStringDiff and significantly different miRNA identified (logPV > 1 & absFC > .58).

### Data analyses and integration

All analyses were undertaken using the R platform for statistical computing (version 3.6.1) and the indicated library packages implemented in Bioconductor.

## Declarations

### Ethics approval and consent to participate

The serum samples from the Southwest Oncology Group (SWOG) clinical trial (SWOG S9917) were collected under local IRB approval [60]. The PCa samples from EA and AA patients who received vitamin D_3_ prior to radical prostatectomy were collected under local IRB approval [7]. The radical prostatectomy PCa samples from EA and AA patients prior were collected under local IRB approval[61]

### Consent for publication

Not applicable

### Availability of data and materials

The datasets generated and/or analysed during the current study will be available on GEO

### Authors’ contributions

*MS* undertook ChIP-Seq, RNA-Seq and cell biology; *SS* undertook ATAC-Seq, RIME, cell biology; *SAW* contributed to ChIP-Seq, ATAC-Seq and RIME; *HT* contributed to bioinformatic analyses; *JSG* contributed to ATAC-Seq and cell biology; *HJ*, *HW* contributed to cell biology; *MDL* undertook GIGGLE analyses; *IE* undertook ancestry informative analyses; *BK*, *HW* contributed to experimental design; *RM*, *GH* undertook the RNA-Seq of the vitamin D supplementation trial in prostate cancer patients; *IBA*, *SOR* contributed to experimental design and conceptual framework; *ARM*, *MBD* and *RAK* designed and oversaw the RNA-Seq of the tumors from prostate cancer patients and captured the clinical data including serum 25(OH)D_3_ levels; *LN* contributed to data interpretation; *JRM* was a co-lead on SWOG S9917 and supplied serum samples for analyses; *CHH* and *GH* oversaw and designed the vitamin D supplementation trial in prostate cancer; *LES-C*, *CY* and *MJC* jointly conceived of the study design; *MJC* oversaw the implementation of the study and undertook other bioinformatic analyses and generated tables and figures.

---

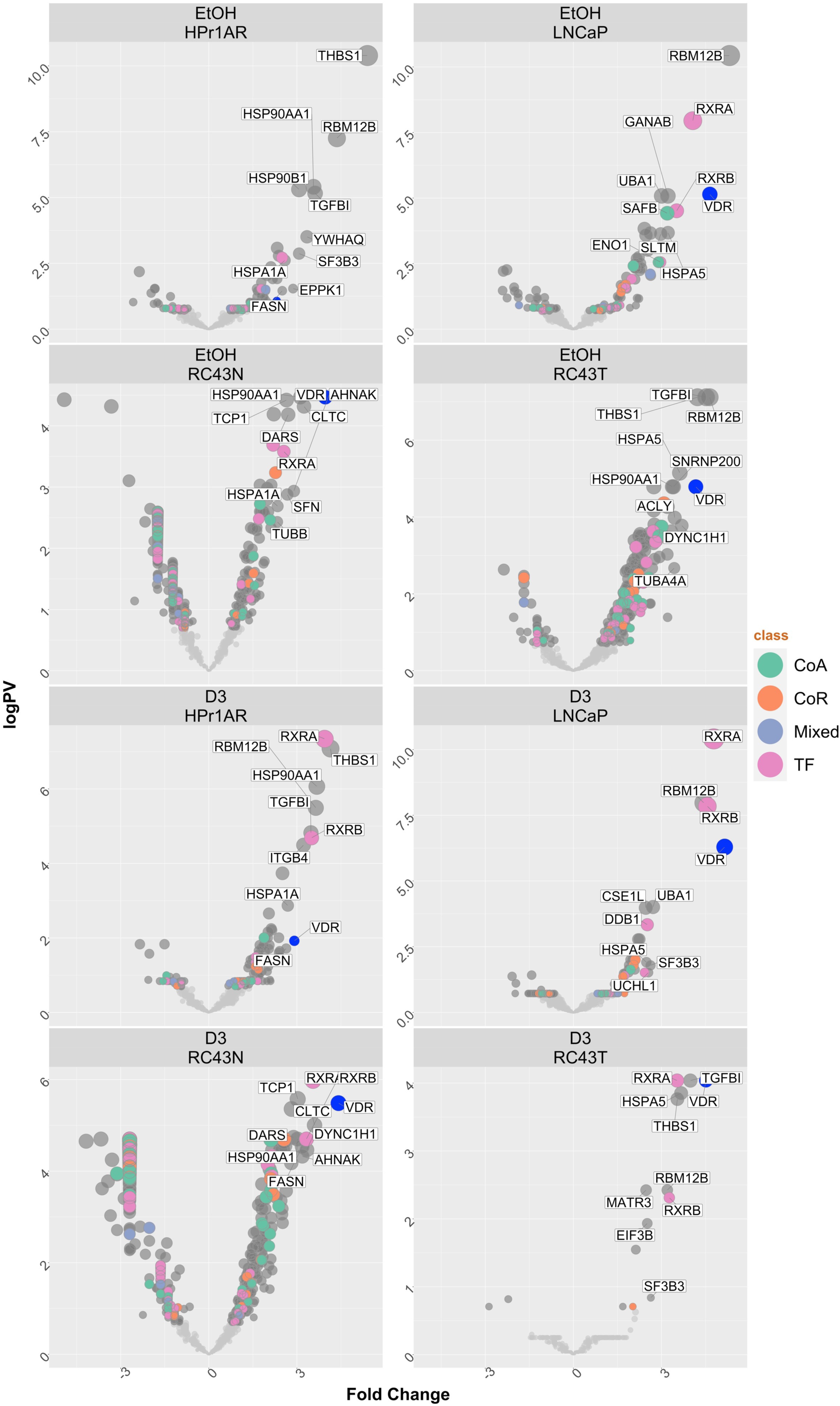

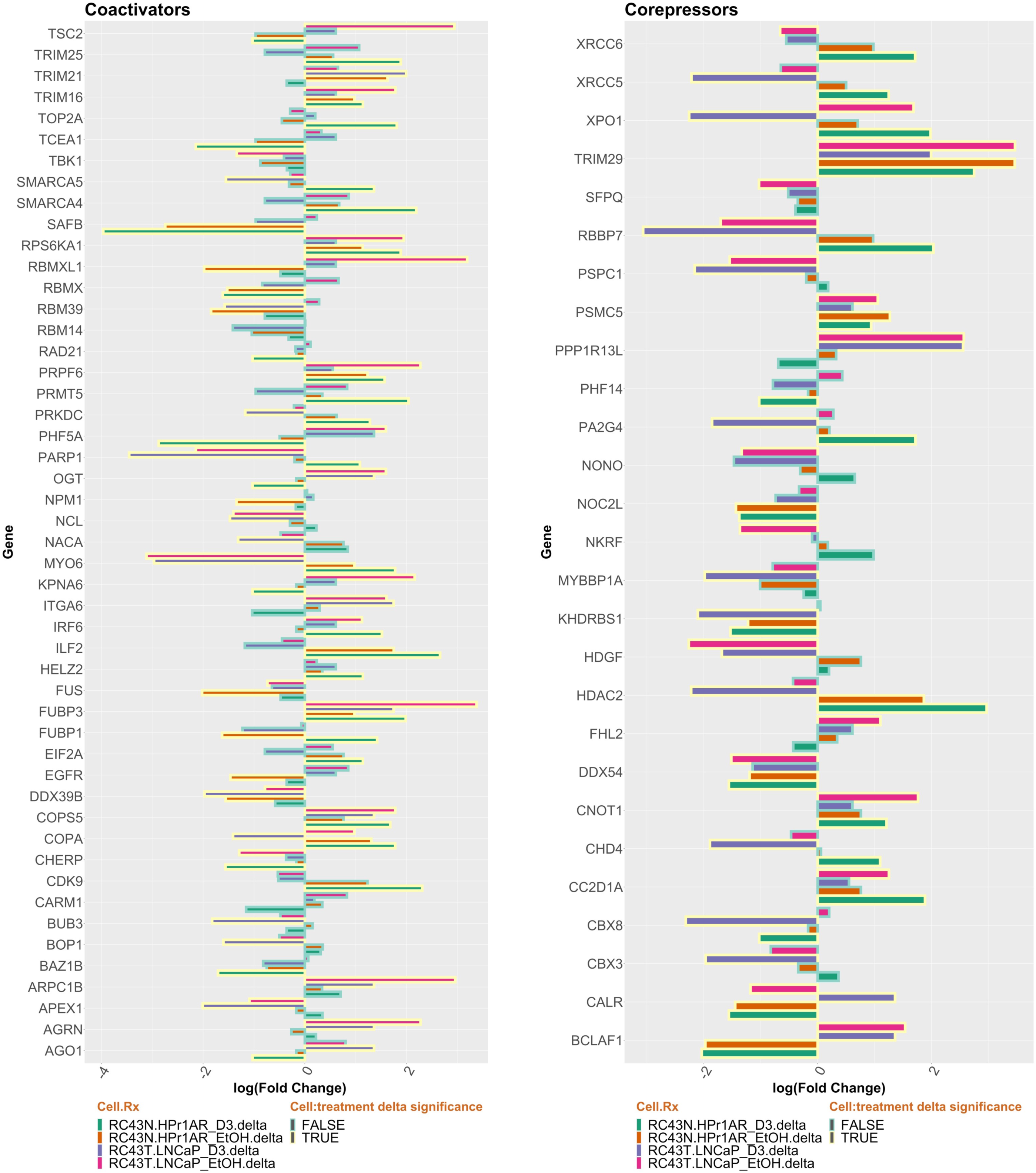

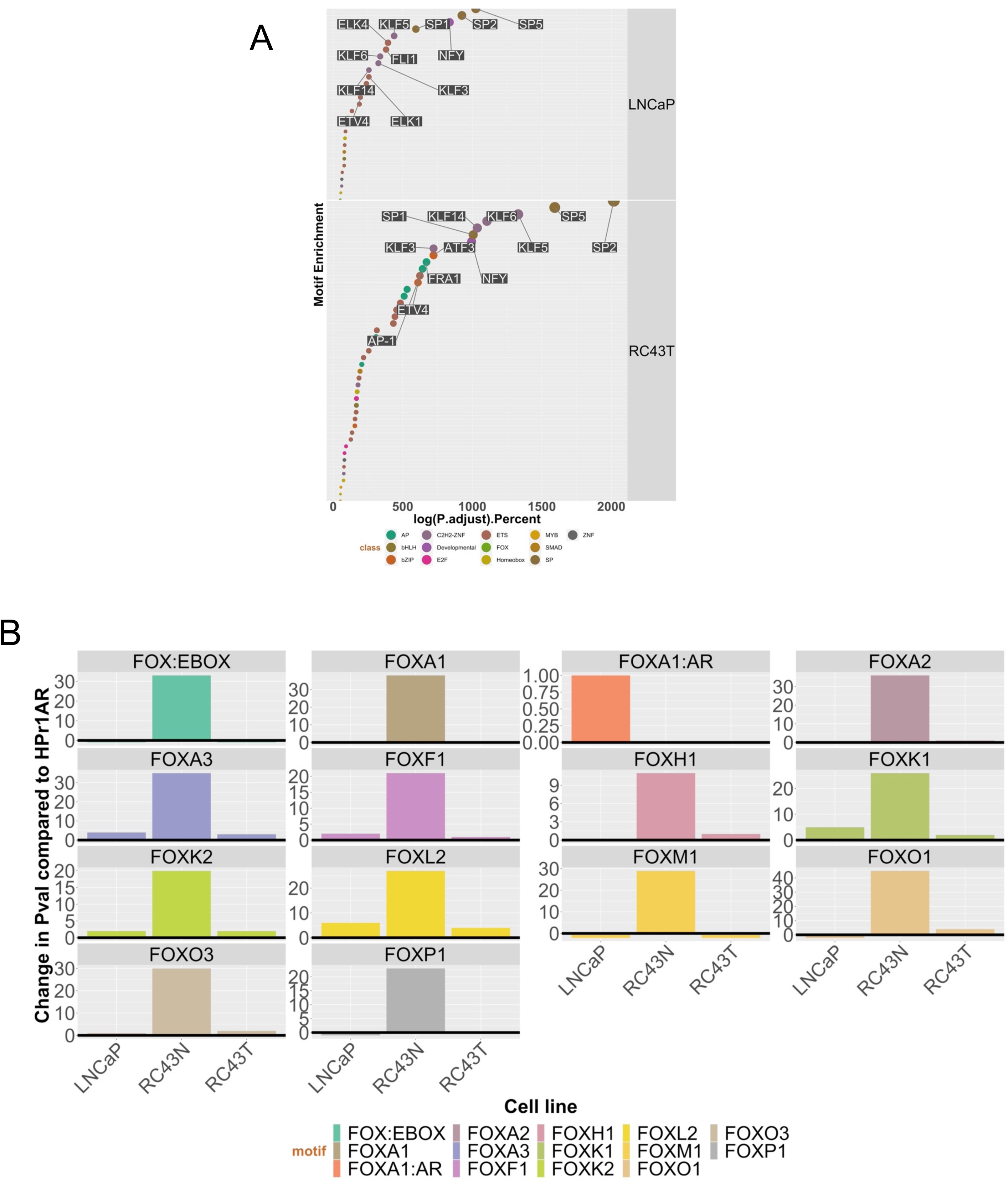

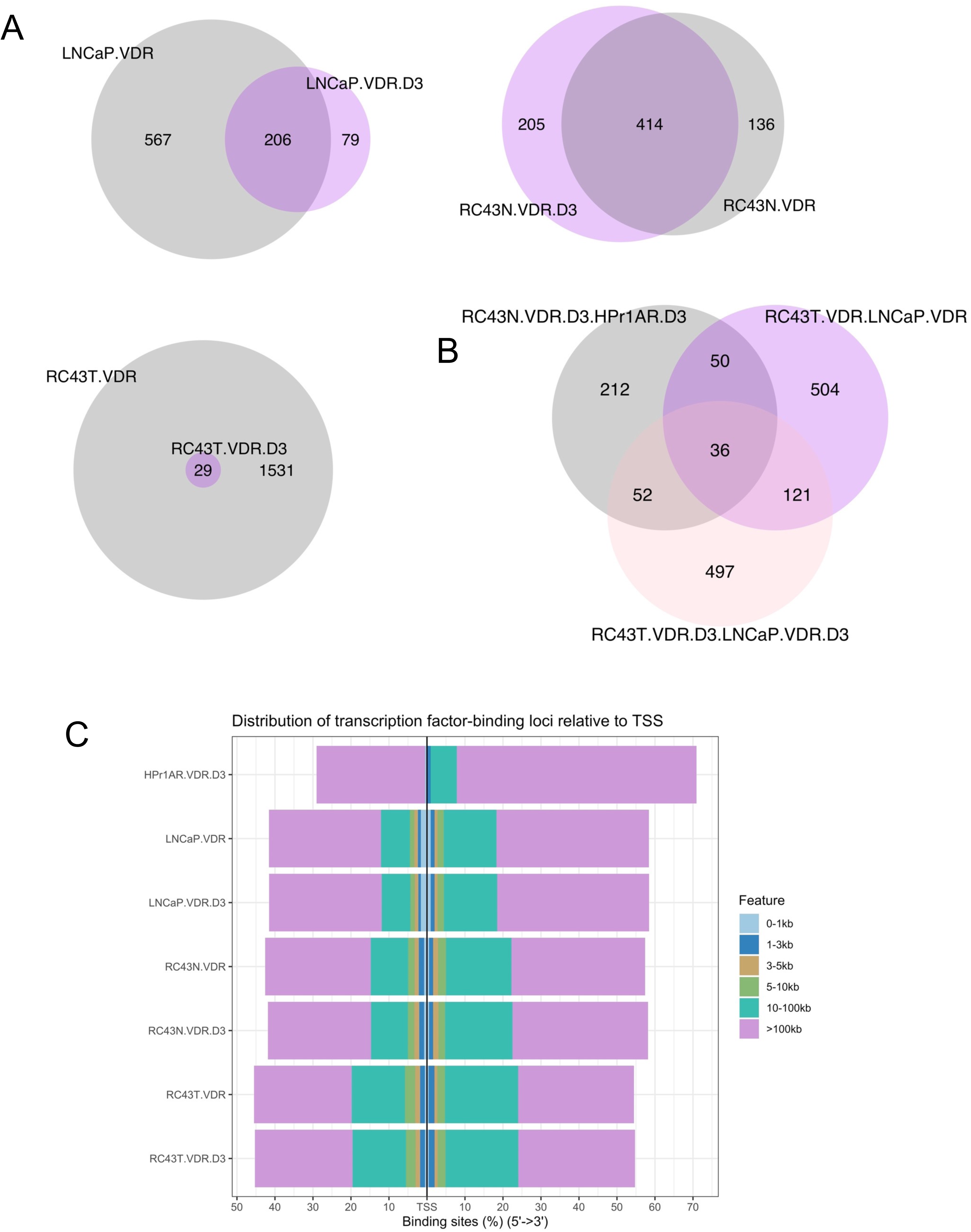

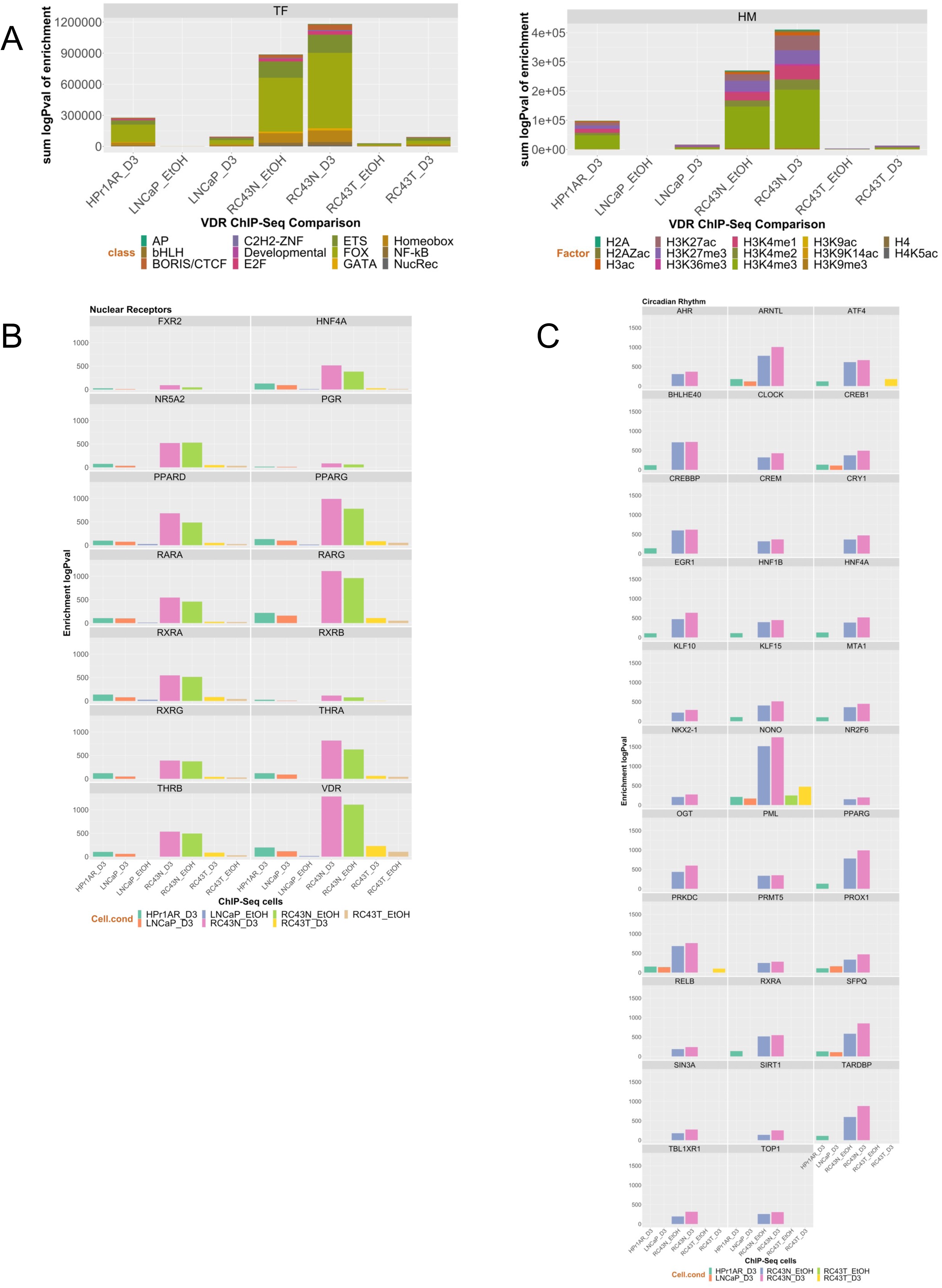

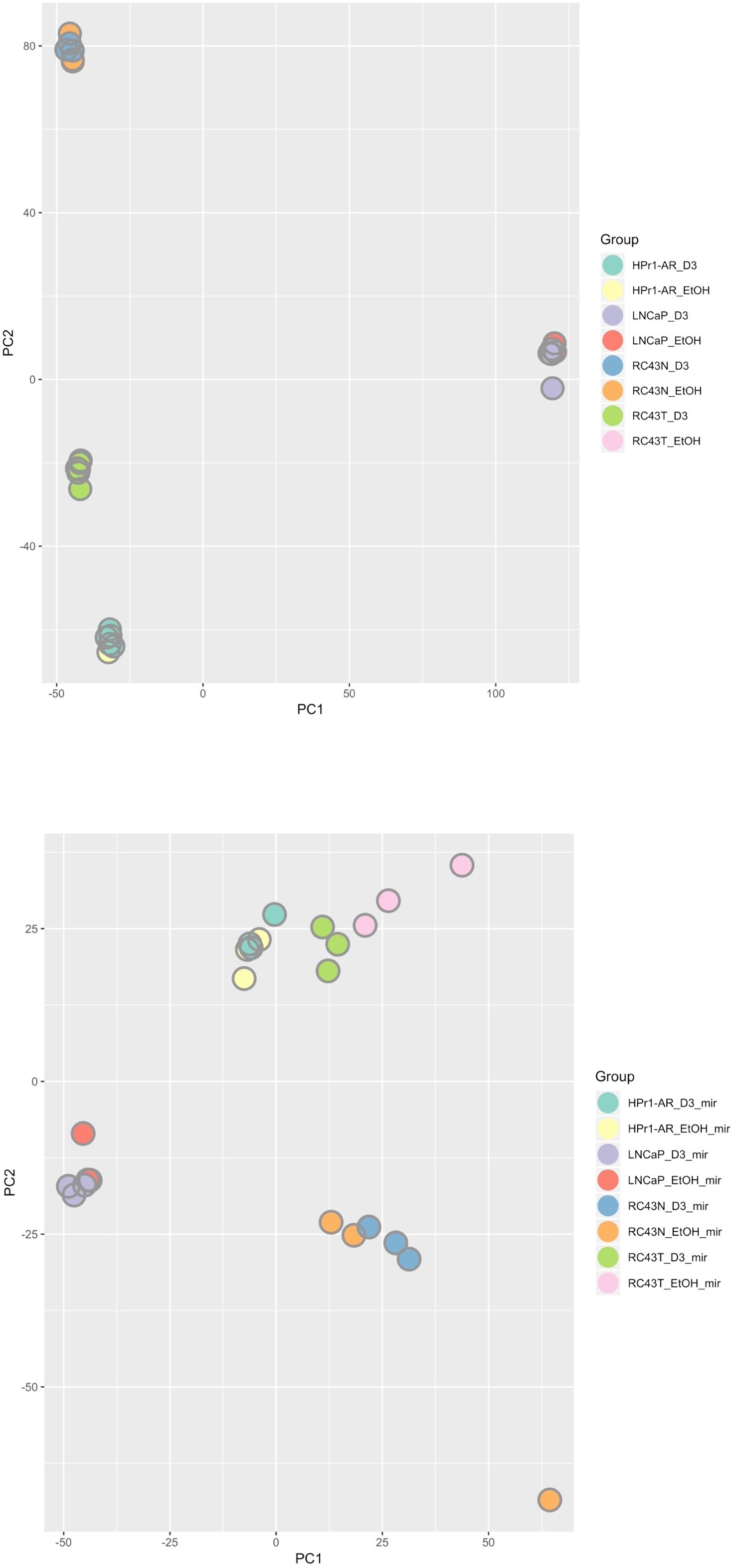

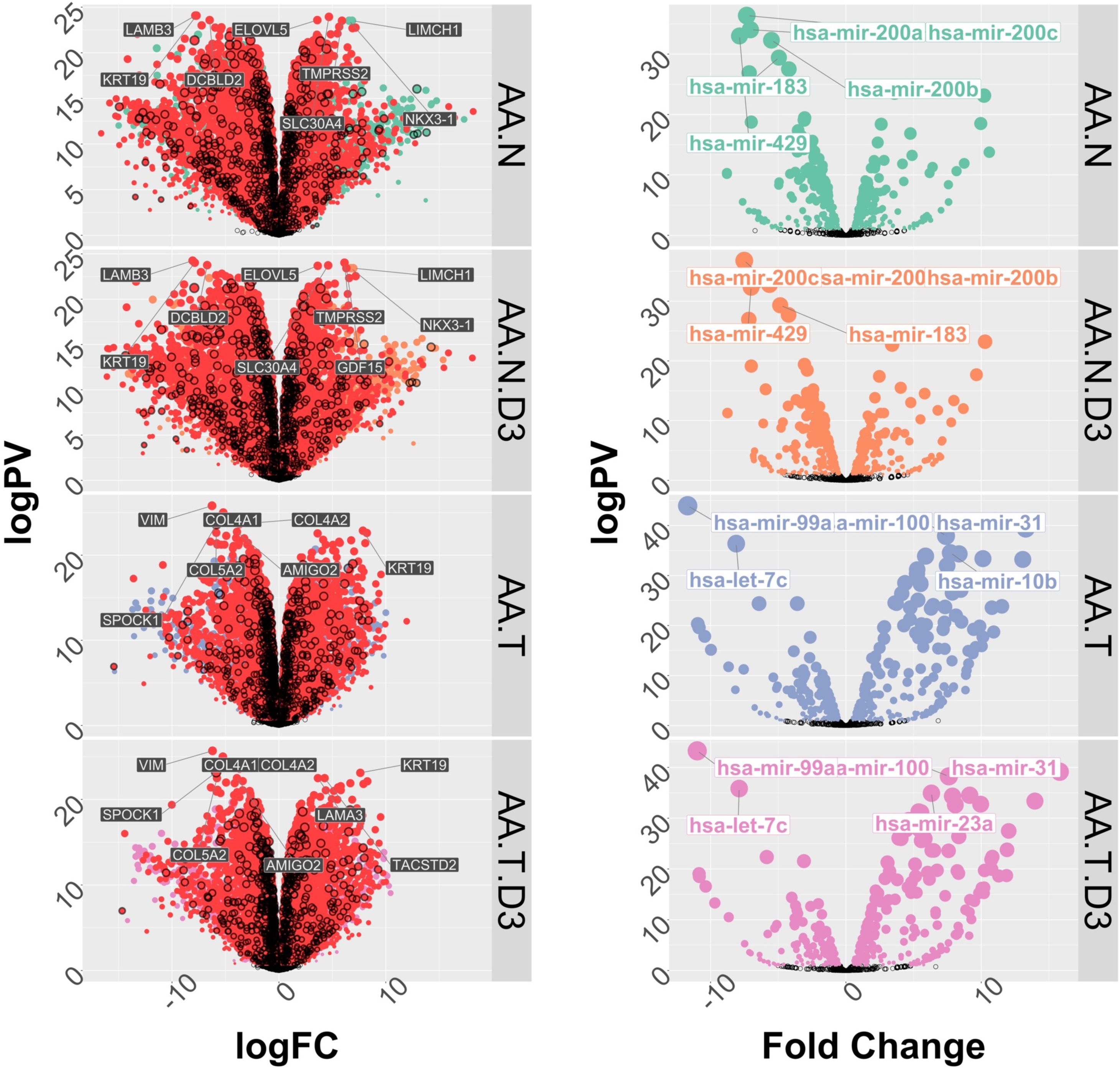

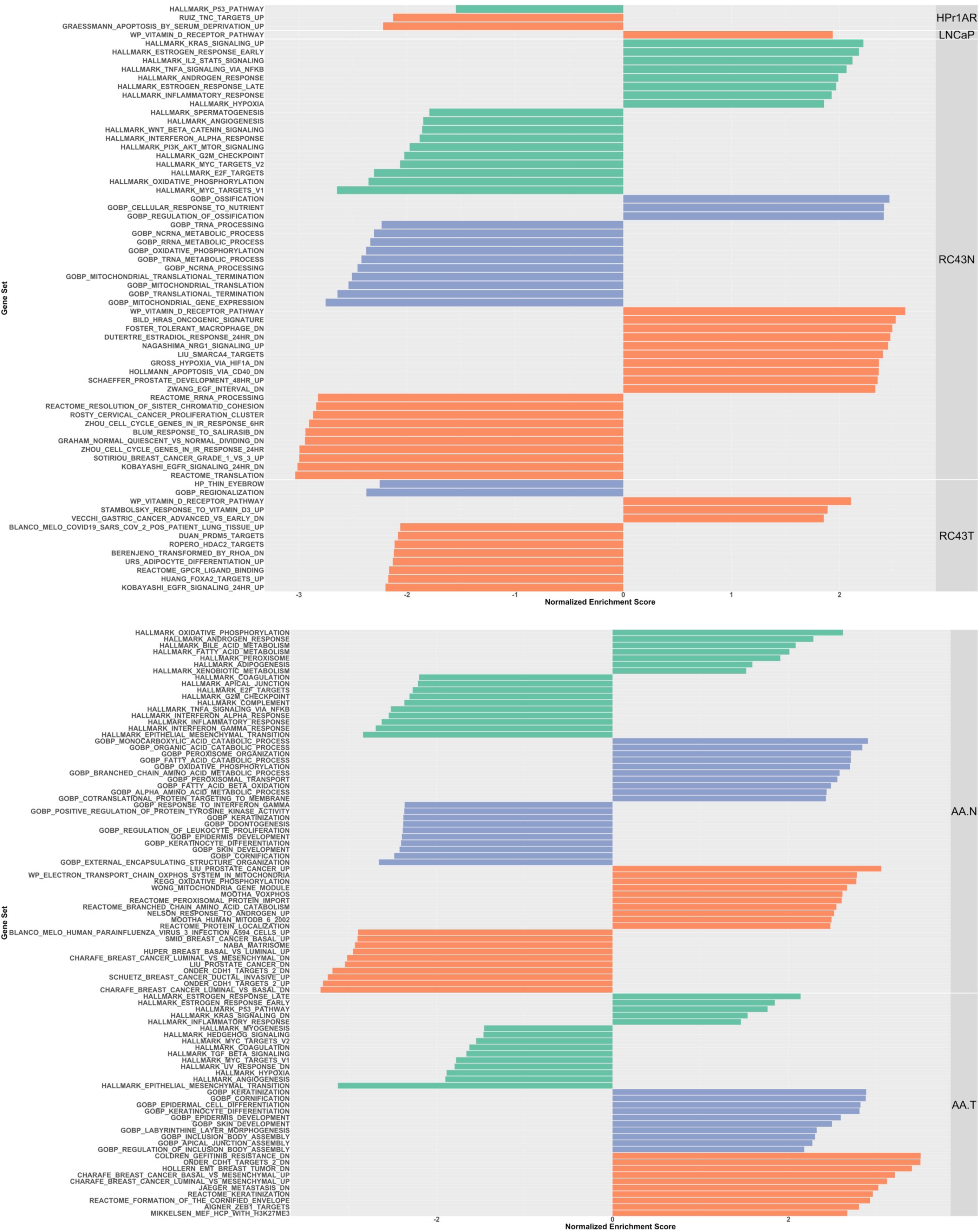

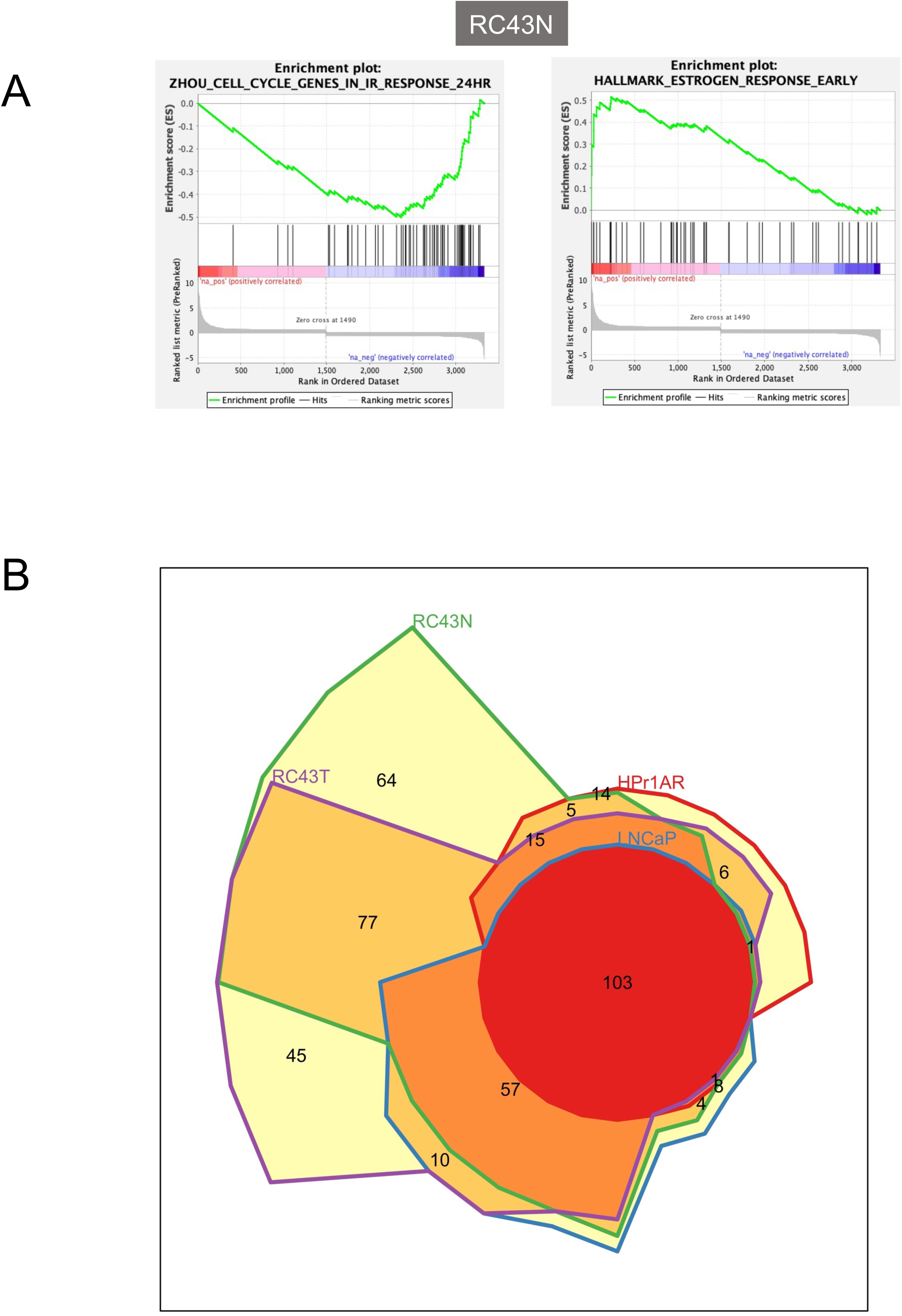

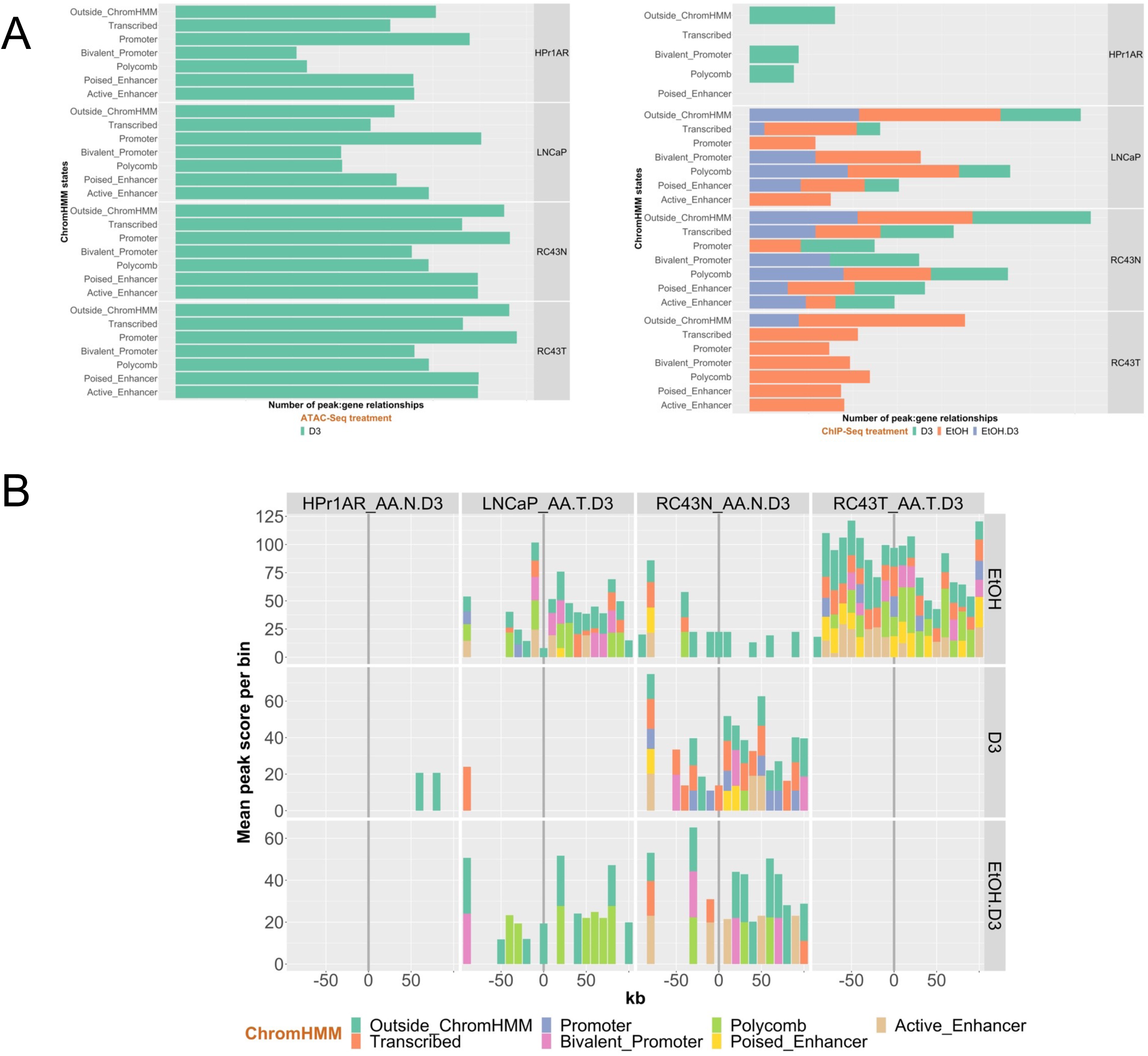

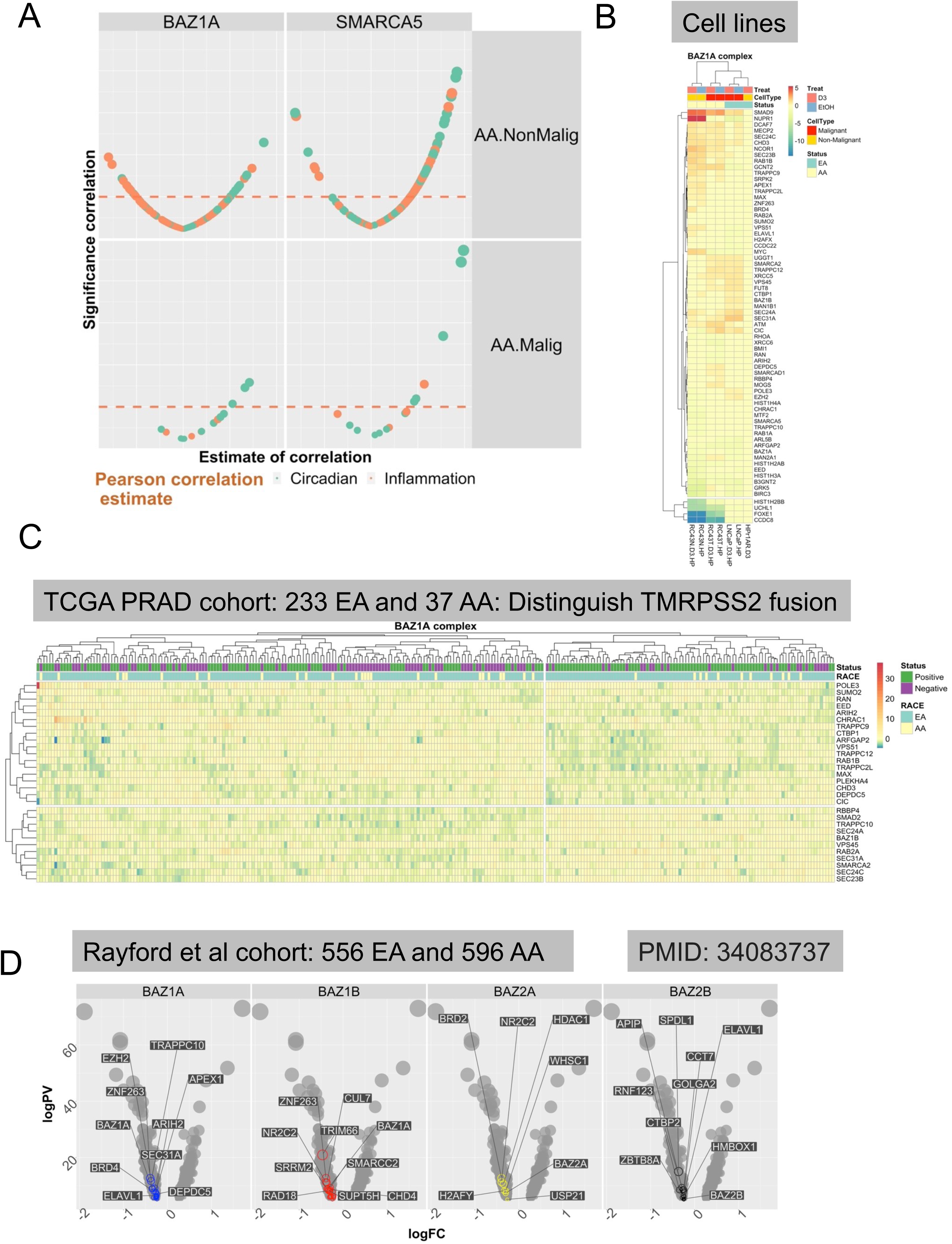

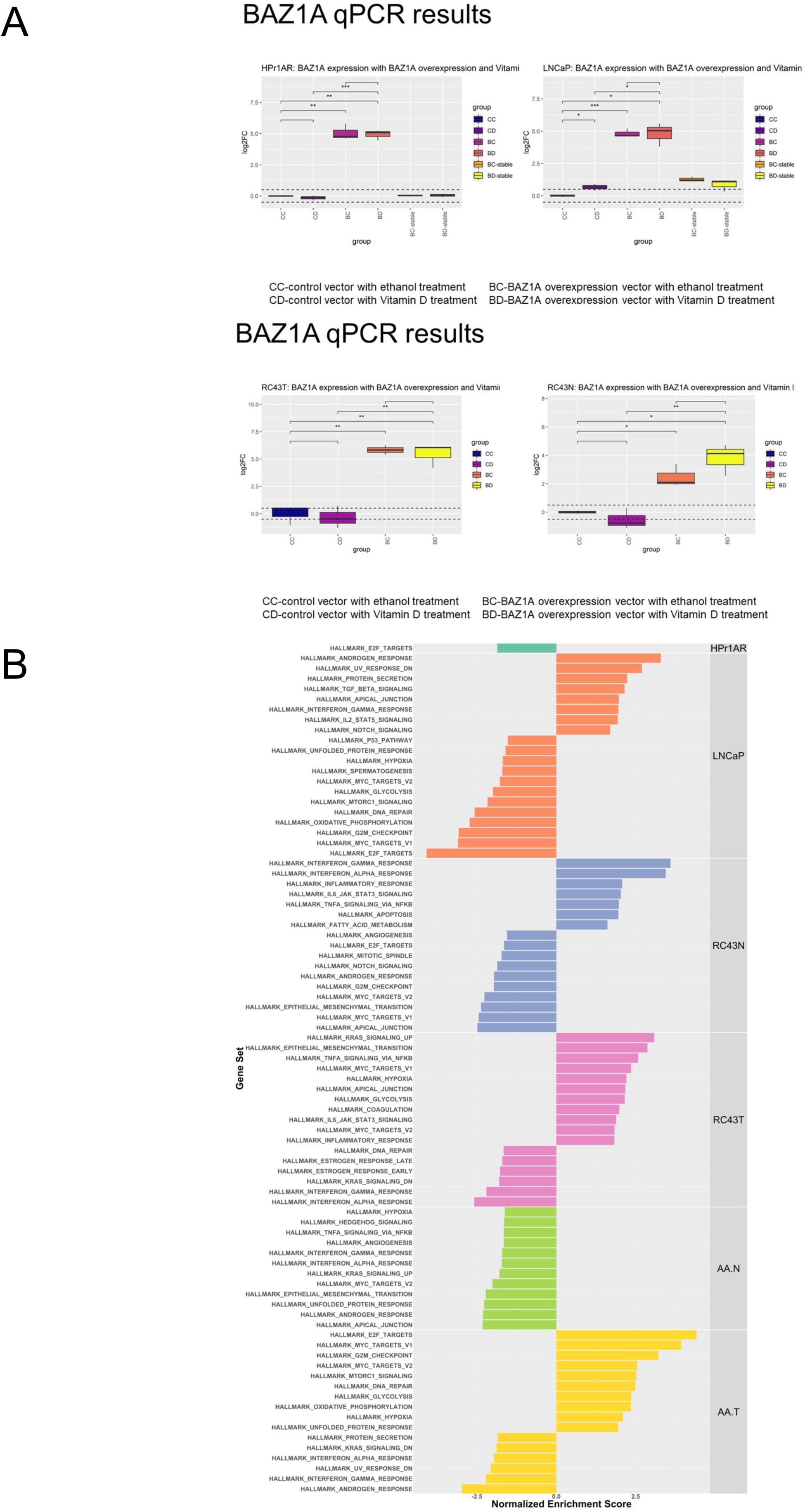

